# Microbial stowaways – waterbirds as dispersal vectors of aquatic pro- and microeukaryotic communities

**DOI:** 10.1101/2021.10.21.465236

**Authors:** Beáta Szabó, Attila Szabó, Csaba F. Vad, Emil Boros, Dunja Lukić, Robert Ptacnik, Zsuzsanna Márton, Zsófia Horváth

**Author notes:** Correspondence Beáta Szabó, Institute of Aquatic Ecology, Centre for Ecological Research, Budapest, Hungary.

## Abstract

**Aim:** Waterbirds are important dispersal vectors of multicellular organisms such as macrophytes, aquatic macroinvertebrates, and zooplankton. However, no study to date has focused on their potential role in dispersing aquatic microbial communities (i.a., bacteria, algae, protozoa). Here, we explicitly studied passive transport (endozoochory) of prokaryotes and unicellular microeukaryotes by waterbirds based on DNA metabarcoding approaches. By directly comparing the dispersed set of organisms to the source pool of a natural metacommunity, we aimed for a realistic estimate of the overall importance of waterbird zoochory for natural microbial communities.

**Location:** Shallow saline temporary ponds (soda pans) in the cross-border region of Austria and Hungary.

**Taxon:** Prokaryotes and unicellular microeukaryotes.

**Methods:** In 2017 and 2018, water samples from 25 natural aquatic habitats along with fresh droppings of the dominant greylag goose (*Anser anser*) and four other waterbird species were collected in a habitat network of temporary ponds. Their prokaryotic and microeukaryotic communities were identified via 16S and 18S rRNA gene amplicon sequencing. Sequence reads were analysed using mothur. After quality filtering of the reads, pro- and microeukaryotic amplicon sequencing variant (ASV) compositions were compared between the aquatic and dropping samples, across years and waterbird species.

**Results:** We found that 28% of the dominant aquatic prokaryotic and 19% of the microeukaryotic ASVs were transported by *A. anser*. ASV richness in *A. anser* droppings was lower, but compositional variation was higher compared to the aquatic communities, probably resulting from stochastic pick-up of microbes from multiple aquatic habitats. We furthermore found that the composition of prokaryotic ASVs in bird droppings were different among the two years and reflected the actual aquatic communities. The dispersed set of microbes were largely similar among the different waterbird species except for the planktivore filter-feeder northern shoveler (*Spatula clypeata*) which was outstanding by dispersing a more species-rich subset of microeukaryotes than shorebirds or geese.

**Main conclusions:** By using a combined amplicon-sequencing approach to characterize microorganisms in waterbird droppings and in the associated environment, our study provides strong evidence for endozoochory of natural aquatic microorganism communities. These results imply that waterbirds may be crucial in maintaining ecological connectivity between discrete aquatic habitats at the level of microbial communities.

## Introduction

Dispersal is a key process connecting habitats, thereby sustaining gene flow (Clobert et al., 2012), biodiversity (Leibold et al., 2004), and ecosystem functions (Bannar-Martin et al., 2018; Zobel et al., 2006). For a long time, prokaryotes, together with unicellular and small multicellular eukaryotes have been considered to have a cosmopolitan distribution and their communities were assumed to be driven only by local environmental and biotic factors (Baas-Becking, 1934; Beijerinck, 1913). However, recent studies (e.g. Cho & Tiedje, 2000; Martiny et al., 2006; Telford et al., 2006; Zinger et al., 2014) benefiting from the rapid development of community sequencing methods led to a paradigm shift by providing evidence for biogeographical patterns and increased recognition of the importance of spatial processes in microorganisms (Langenheder & Lindström, 2019; Mony et al., 2020; Ptacnik et al., 2010; van der Gast, 2015; Vyverman et al., 2007). This has finally placed microbes in the same metacommunity framework that has been already well-established for macroorganisms (Leibold & Chase, 2018).

Hence, the importance of passive dispersal for microorganisms is now acknowledged, which can occur by wind (Genitsaris et al., 2011; Sharma et al., 2007), water currents (Luef et al., 2007), animals (Figuerola & Green, 2002a; Green et al., 2008; Valls et al., 2017), and human activities (Reise et al., 1999; Ruiz et al., 2000). But despite the increasing interest in microbial dispersal and the availability of modern molecular techniques, zoochory is still largely neglected in this respect. Although there is evidence for waterbirds being effective short- and long-distance dispersal agents of macrophytes, macroinvertebrates, zooplankton and vertebrates (Brochet, Gauthier-Clerc, et al., 2010; Figuerola & Green, 2002b; Figuerola et al., 2003; Lovas-Kiss et al., 2019, 2020; Reynolds & Cumming, 2016; Silva et al., 2019; Viana et al., 2013a, 2013b), waterbird-mediated dispersal of unicellular microorganisms (especially bacteria) is poorly understood. There is evidence for the transport of viruses (Blagodatski et al., 2021) and microorganisms exemplified mainly by the dispersal of single and/or pathogenic microbial taxa (Briscoe et al., 2021; Garmyn et al., 2012; Hartikainen et al., 2016; Jarma et al., 2021; Lewis et al., 2014) or their co-dispersal with their infected hosts (Okamura et al., 2019). However, no explicit studies have so far targeted the dispersal potential of waterbirds for natural aquatic microbial communities with a direct comparison of natural communities to taxa dispersed by waterbirds.

Here, we carry out an extensive study on the role of waterbirds as dispersal agents of aquatic pro- and eukaryotic unicellular microorganisms with the help of high-throughput DNA sequencing. Our study area is a landscape of saline temporary ponds, representing a well-delineated habitat network. The characteristic species of the waterbird community in the area is greylag goose (*Anser anser*), with more than 6,000 individuals (Wendelin & Dvorak, 2020). This species is known to be a regular large-bodied visitor of aquatic habitats, moving in flocks of up to 750 individuals (McKay et al., 2006). It has been suggested that they may contribute significantly to the transport of passively dispersing organisms across aquatic habitats (García-Álvarez et al., 2015; Green et al., 2002). However, we lack empirical data to assess their actual role as dispersal agents for microbial organisms.

In line with this, our main objective is to investigate the potential of zoochory by waterbirds for dispersing microorganisms among local habitats in a metacommunity. Specifically, our first aim is to reveal what proportion of the amplicon sequence variants (ASVs) occurring in the aquatic habitats can be found in droppings of the dominant waterbird of the region, *A. anser*. Here, we also investigate whether the microbial communities detected in the bird droppings reflect a possible change of the communities in the aquatic habitats over time. And finally, we assess the dispersal potential of *A. anser* relative to three other waterbird species with different feeding habits and habitat use in the same landscape.

## Material & methods

### Sampling and sample processing

The study area (∼ 200 km^2^, Horváth et al., 2016) in the cross-border region of Fertö / Neusiedlersee Cultural Landscape in Austria and Hungary is characterized by a dense cluster of temporary saline ponds (soda pans). These habitats form a habitat network relatively isolated from freshwater habitats or other soda pans in the central and eastern regions of Hungary (Tóth et al., 2014). The clumped nature of this pondscape, with shallow (≤ 1 m) and hypertrophic aquatic habitats (Boros et al., 2017) offers excellent feeding grounds for invertivorous waterbirds (Horváth et al., 2013) and breeding sites for several other species, including greylag geese (*Anser anser*, Dvorak et al., 2020; Wendelin & Dvorak, 2020). The region is legally protected as part of two national parks (Neusiedlersee-Seewinkel in Austria, and Fertő-Hanság in Hungary), designated as Important Bird Area (BirdLife International, 2021a, 2021b) and part of a UNESCO World Heritage site (Fertő / Neusiedlersee Cultural Landscape).

We collected water samples from 25 soda pans in two consecutive years (3-6 April 2017 and 2-4 April 2018; Figure 1), representing all habitats that held water in both years (hereafter aquatic community samples). The sampled habitats are situated within 17 km (largest distance between two habitats), thereby representing a region where waterbirds can regularly move around on a daily basis (Bell, 1988, Boos et al., 2019; Link et al., 2011; Nilsson & Persson, 1992). From each soda pan, a total of 20 L of water was collected from 20 different points using a one-litre plastic beaker (thus collecting a pooled sample from the largest possible area) and sieved through a 100-μm mesh plankton net to remove large zooplankton and filamentous algae which would hinder the detection of unicellular organisms during amplicon sequencing. Sampling of water was carried out by wading so that we gently collected water from the undisturbed areas in front of us. For further processing, 1 L of the composite sieved water was immediately delivered to the laboratory in a glass bottle in a cool box. As many of the studied soda pans have high turbidity and high prokaryotic or algal cell numbers (10C-10C cells mL^-1^, Boros et al., 2017; Kirschner et al., 2002), for molecular analysis, 1-50 mL of water (depending on turbidity, as Secchi depth ranged from 0.3 to 44 cm) was filtered through a nitrocellulose membrane filter (Ø 47 mm) with a pore size of 0.22 μm until clogging. Thereafter, filters were stored at -20 °C until DNA extraction.

**Figure 1.**
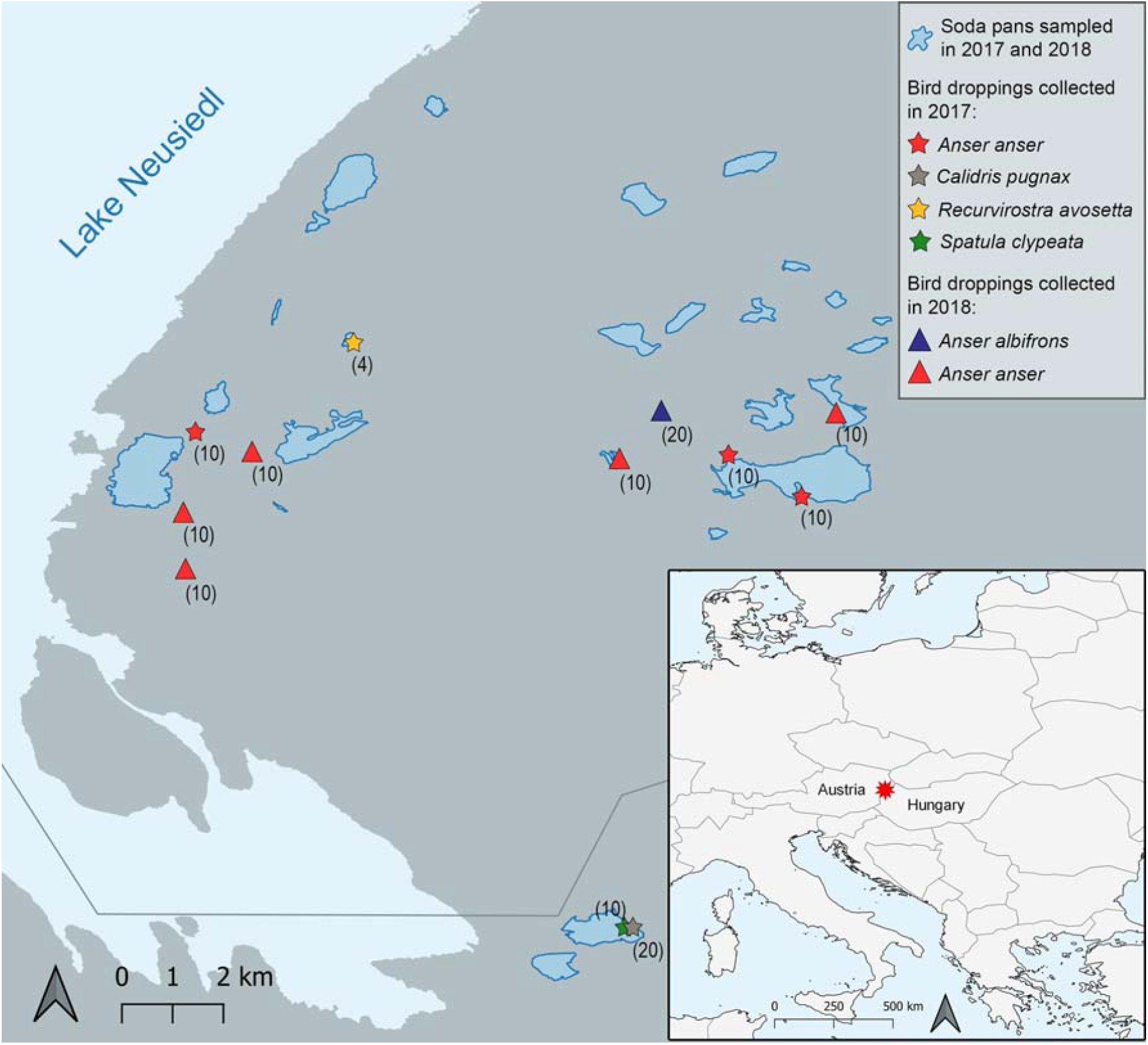
Map of the region and the sampling sites. Projection of the map is EPSG:4326 (WGS84). All the soda pans from which samples were collected are indicated with blue polygons, while symbols highlight the spots where bird dropping samples were collected. The number of collected bird dropping samples is indicated in brackets.

Simultaneously, we collected fresh waterbird droppings at all sites that hosted a monospecific flock of waterbirds. We approached the birds roosting on dry mudflats or grasslands on the shores or right next to the soda pans and once they took off, fresh droppings were collected in sterile cryogenic vials and immediately frozen on dry ice. Droppings at least one-metre apart were collected thereby ensuring an individual being sampled only once (Lovas-Kiss et al., 2018). To scrape off any soil or plant material and pick up the faecal sample we used the vial and its cap, in which the given sample was stored. This way we avoided the potential contamination due to long-term exposure to e.g. wind-dispersed propagules and also avoided possible cross-contamination with a shared sampling equipment. Bird droppings were stored at -20 °C until further processing. In 2017, a total of 64 droppings from *Anser anser, Calidris pugnax, Recurvirostra avosetta, Spatula clypeata*, while in 2018, altogether 70 droppings from *A. anser* and *A. albifrons* were collected with this method (Figure 1). The feeding mode of the waterbird species was summarized in Table S1.

We used the two datasets (aquatic communities and bird droppings) to compare the possible dispersal provided by waterbirds with the implicit limitation that we cannot differentiate between viable propagules and non-viable remnants of the original microorganisms. At the moment, there is no single culturing method that could have been applied for waterbird droppings without being extremely selective for the emerging microbes and hence we decided to sequence the samples as a whole (as in Hartikainen et al., 2016 and Jarma et al., 2021). Although this can mean an overestimate for the ratio of successfully dispersed taxa, it can still provide a critical first estimate of what might be transported by the birds, especially given by their inefficient digestion (Frisch et al., 2007; Green & Sánchez, 2006; Lovas-Kiss et al., 2020) and short retention times (Brochet, Guillemain, Gauthier-Clerc et al., 2010; Sánchez et al., 2012).

### DNA isolation, amplification and sequencing

DNA extraction from the filters and waterbird droppings was performed after the sampling campaign in 2018. The PowerSoil® DNA Isolation Kit (MO BIO Laboratories Inc., Carlsbad, CA, USA) was used, following the manufacturer’s instructions. Extracted DNA samples were stored at -20 °C before shipping for amplification and sequencing at an external company (LGC Genomics, Berlin).

Prokaryotic 16S rRNA and microeukaryotic 18S rRNA gene amplification was carried out using the following primer pairs: EMBf 515F (GTGYCAGCMGCCGCGGTAA, Parada et al., 2016) – EMBr 806R (GGACTACNVGGGTWTCTAAT, Apprill et al., 2015) for the V4 region of the prokaryotic 16S rRNA gene and UnivF-1183mod (AATTTGACTCAACRCGGG) – UnivR-1443mod (GRGCATCACAGACCTG) (Ray et al., 2016) for the V7 region of the eukaryotic 18S rRNA gene. For each sample, the forward and reverse primers had the same 10-nt barcode sequence. Amplification of the target taxonomic marker gene regions and sequencing were carried out by LGC Genomics (Berlin, Germany). The PCRs included about 1-10 ng of DNA extract (total volume 1μl), 15 pmol of each forward primer and reverse primer in 20 μL volume of 1 x MyTaq buffer containing 1.5 units MyTaq DNA polymerase (Bioline GmbH, Luckenwalde, Germany) and 2 μl of BioStabII PCR Enhancer (Sigma-Aldrich Co.). Amplification was carried out for 30 cycles using the following parameters: 1 min 96 °C pre-denaturation; 96 °C denaturation for 15 s, 55 °C annealing for 30 s, 70 °C extension for 90 s, hold at 8 °C. About 20 ng amplicon DNA of each sample were pooled for up to 48 samples carrying different barcodes. The amplicon pools were purified with one volume Agencourt AMPure XP beads (Beckman Coulter, Inc., IN, USA) to remove primer dimers and other small mispriming products, followed by an additional purification on MiniElute columns (QIAGEN GmbH, Hilden, Germany). About 100 ng of each purified amplicon pool DNA was used to construct Illumina libraries using the Ovation Rapid DR Multiplex System 1-96 (NuGEN Technologies, Inc., CA, USA). Illumina libraries (Illumina, Inc., CA, USA) were pooled and size selected by preparative gel electrophoresis. Sequencing was performed on an Illumina MiSeq using 300 bp paired-end format with a V3 Reagent Cartridge on the Illumina MiSeq platform, aiming for 50,000 raw sequence read pairs per sample. Sequencing was carried out in 48 samples/plate or 24 samples/plate format to decrease overall index-hopping bias between samples. Only five lake samples were sequenced at the same time with bird dropping samples in 2 out of 13 runs.

### Amplicon data analysis

Sequence processing, taxonomic assignments and ASV picking were carried out with mothur v1.43.0 (Schloss et al., 2009) using the MiSeq SOP as reference (http://www.mothur.org/wiki/MiSeq_SOP; Kozich et al., 2013, downloaded on 12^th^ November 2020). Additional quality filtering steps were also applied to eliminate possible sequence artifacts, such as the adjustment of the deltaq parameter to 10 in the ‘make.contigs’ command, primer removal from both ends of the sequences and the exclusion of singleton reads according to Kunin et al. (2010) before ASV identification. Denoising was performed using mothur’s pre.cluster command using the default algorithm and applying the suggested 2 bp difference cutoff. Chimeras were identified and removed using the mothur implemented version of VSEARCH. Read alignment and taxonomic assignment were carried out using the ARB-SILVA SSU Ref NR 138 reference database with a minimum bootstrap confidence score of 80 (Quast et al., 2013). ASVs assigned to non-primer-specific taxonomic groups (‘Chloroplast’, ‘Mitochondria’ and ‘unknown’) were subsequently removed from the dataset. For prokaryotic ASVs, the TaxAss software (Rohwer et al., 2018) was used with default parameters for taxonomic assignment with the FreshTrain (15 June 2020 release) and ARB-SILVA SSU Ref NR 138 databases, while taxonomic assignment of the 18S rRNA gene ASVs was performed using the PR2 v4.12.0 reference database (Guillou et al., 2013).

ASVs assigned to taxa Streptophyta, Metazoa, Ascomycota and Basidiomycota were excluded from the eukaryotic ASV set. Subsequently, both 16S and 18S ASV sets were rarefied to the read number of the sample having the lowest sequence count (8620 read per sample for the 16S set and 2432 read per sample for the 18S set).

### Statistical analysis

We used the rarefied 16S (hereinafter referred to as prokaryotes) and 18S (microeukaryotes) community datasets separately in all our analyses. As our main aim was to quantify the potential dispersal of aquatic microorganisms by waterbirds, we excluded those organisms (both prokaryotes and microeukaryotes) that are likely not members of the natural aquatic community of the water samples. Accordingly, in case of both the aquatic community samples and the bird droppings, we only used ASVs that were present at least in one aquatic community sample with ≥1% relative abundance (“aquatic subset”). Since the subsetting resulted in different read numbers per sample, we converted the abundances to relative abundances prior to the statistical analyses. In the main part of the manuscript, we used only these aquatic subsets. This subsetting was carried out separately for prokaryotes and microeukaryotes. The resulting aquatic subset of waterbird droppings contained 1.9% (±4.1%) of the original prokaryotic and 4.5% (±6.8%) of the microeukaryotic ASV abundances in these samples. The subset of aquatic communities contained 71.0% (±12.7%) of the original prokaryotic and 84.2% (±9.1%) of the microeukaryotic ASV abundances detected in the unselected datasets. The unselected ASV sets, the aquatic subsets and the related list of taxa were presented for both prokaryotes and microeukaryotes as supplementary data (Table S2-S9).

For a quantitative assessment of waterbird dispersal potential in the pondscape, we only used *A. anser* samples, being the only species from which we could collect samples in both years. To exclude a potential bias arising from the different sampling effort in soda pans vs bird droppings, a random re-sampling was performed based on the lowest sample size for both prokaryotes (n=19) and microeukaryotes (n=9) per sample group, resulting in a total of 76 pro- and 36 microeukaryote samples used in these comparisons.

To estimate the possible significant effect of sample type (*A. anser* droppings vs. aquatic communities) and sampling year (2017 vs. 2018) on the local ASV richness (α-diversity) and compositional change among samples (Whittaker’s β-diversity: β = γ/α), non-parametric Scheirer-Ray-Hare test with an interaction term was run using the ‘rcompanion’ v. 2.4.6 package (Mangiafico, 2021), followed by Dunn’s post-hoc test for pairwise comparisons with ‘FSA’ v. 0.9.1 package (Ogle et al., 2021) where p-values were adjusted with the Benjamini-Hochberg method.

We created stacked barplots to illustrate the quantitative differences of the higher-order prokaryotic and eukaryotic taxa among the sample groups. Prior to this, third level taxon names were assigned to the ASVs detected in the samples, thereafter ASV abundances belonging to the same taxon were summed up and expressed as relative abundance in each sample group. Taxa that did not reach 4% relative abundance at least in one of the four sample groups were combined in the category “Other”.

Principal coordinate analysis (PCoA) was performed to illustrate the separation of samples according to sample type and sampling year with the ‘vegan’ v. 2.5-7 package (Oksanen et al., 2020). To test for significant differences in the same dataset, two-way PERMANOVA with an interaction term (based on 2000 permutations) was carried out, followed by a pairwise comparison of the four sample groups (based on 2000 permutations) with the ‘pairwiseAdonis’ v. 0.0.1 package (Arbizu, 2017). We ran additional SIMPER analyses to determine which ASVs are the most responsible for the dissimilarities among sample types and sampling years. In order to provide comparable results, PCoA, PERMANOVA, pairwise comparison and SIMPER were all run based on Bray-Curtis dissimilarity calculated from ASV relative abundance data.

We repeated our analyses based on the unselected datasets (i.e. without selecting for aquatic taxa) and presented those results in the Supplementary material (Table S11, S14, Figure S1, S2, S4-S6, S9-S10). To standardize sample sizes, re-sampling was carried out also for the unselected dataset of prokaryotes (n=25) and microeukaryotes (n=10) based on the lowest sample size resulting in a total of 100 pro- and 40 microeukaryote samples.

To compare prokaryotic and microeukaryotic ASV richness in each sample group (droppings of different waterbird species and aquatic community samples from both years), we applied sample-size-based rarefaction and extrapolation approach (Chao et al., 2014) using ‘iNEXT’ v. 2.0.20 package (Hsieh et al., 2020). The 95% confidence intervals were constructed by bootstrapping (based on 50 bootstrap replications).

To reveal if different waterbird species transport different microbial communities, and whether they differ among the two sampling years, we performed separate PCoA analyses including the waterbird species from which samples were collected in at least one year (sample numbers after re-sampling the amplicon data are presented in Table S1).

We excluded *R. avosetta* from the comparative analyses of different waterbird species and aquatic community samples due to the low number of samples (Table S1). However, we present the raw sequence reads in the data depository and the ASV sets with the related taxonomic list as supplementary files (Table S2-S9) for each of the five waterbird species.

All analyses focusing on community composition were furthermore repeated for incidence data based on Sørensen dissimilarity.

Statistical analyses were carried out using R v. 4.1.1 statistical software (R Core Team, 2021).

## Results

We found a consistent difference between the number of prokaryotic and microeukaryotic ASVs in the two main sample types (*Anser anser* droppings and aquatic communities) after rarefaction. Local ASV richness (α-diversity) was significantly higher in the aquatic community samples (in both years), while compositional variation (β-diversity) was higher among the *A. anser* samples, especially in prokaryotes. In line with the local richness, regional ASV richness (γ) was also higher in the aquatic communities in both prokaryotes and microeukaryotes. In general, there was no remarkable difference in the diversity metrics between the two sampling years (Figure 2, Table S10), however, in 2017, microeukaryotic β-diversity did not differ significantly between the two sample types (aquatic community and *A. anser* droppings). When repeating the analyses for the unselected prokaryotic and microeukaryotic community datasets (thereby also including the gut microbiome, possible parasites of waterbirds and other non-aquatic microorganisms), patterns of α- and γ-diversity were similar to the results based on the aquatic subset (i.e. less ASVs in *A. anser* samples independent of sampling year), however, β-diversity was low in case of both sample types in both years (Figure S1, Table S11).

**Figure 2.**
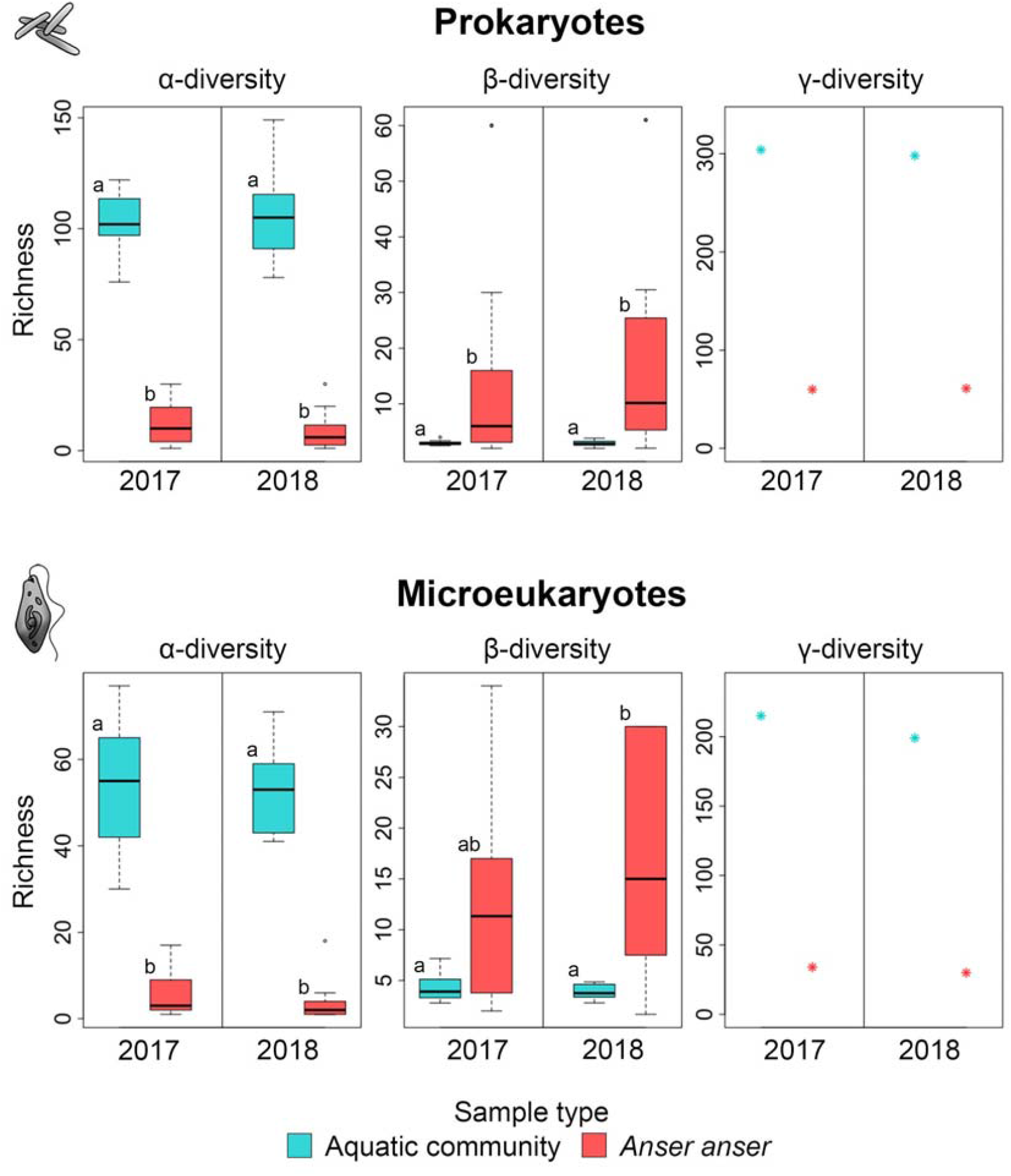
α-, β- and γ-diversity of the prokaryotic and microeukaryotic aquatic subsets in aquatic communities and *Anser anser* droppings in 2017 and 2018. Different letters indicate statistically significant differences in α- and β-diversity at a significance level of p_adj_<0.05 based on Dunn’s pairwise post-hoc test. Pairwise γ-diversity comparisons are presented in Figure 5.

In line with this, the majority of ASVs were found only in the aquatic habitats, with most ASVs shared between years (Figure 3, Venn diagrams). Even so, we detected a considerable proportion of ASVs present in aquatic habitats also in the *A. anser* samples: 28% of the prokaryotic and 19% of the microeukaryotic ASVs were shared among both types of samples, with 19-19% (2017 and 2018, prokaryotes) and 9-10% (2017 and 2018, microeukaryotes) of ASVs being shared among birds and aquatic communities within the same year (Figure 3). Among prokaryotes, 9% of the ASV were found in all four sample groups (both sample types in both years; Figure 3). Compared to this, the share of microeukaryotic ASVs present in all four sample groups was low (3%) (Figure 3). In the unselected community datasets, trends and differences were similar to those observed in our aquatic data subsets, except for the high number of ASVs unique to *A. anser* samples (33% for prokaryotes and 26% for microeukaryotes) (Figure S2).

**Figure 3.**
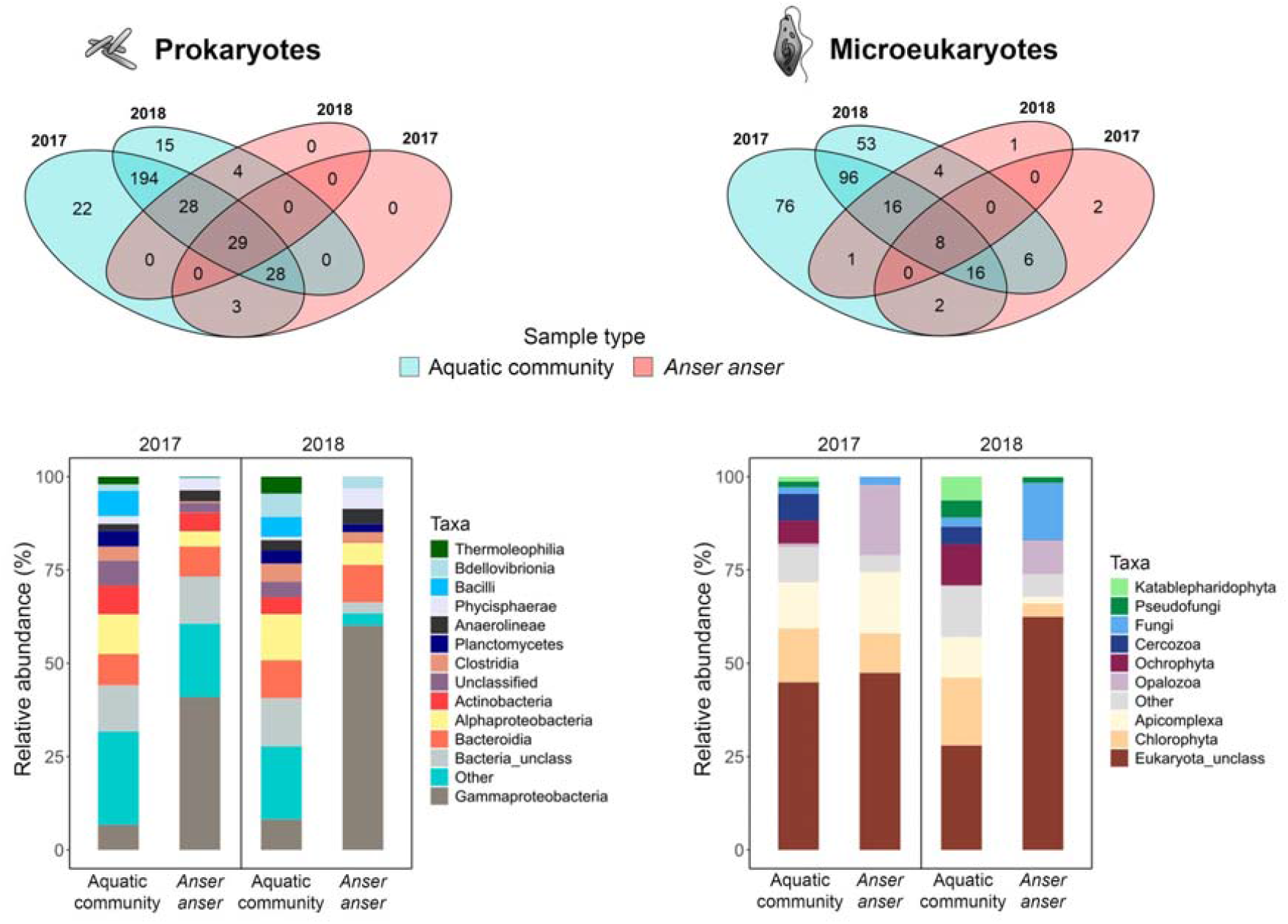
Number of prokaryotic and microeukaryotic ASVs (above) shared among sample types (aquatic community and *Anser anser* dropping) and years, and the relative abundance of higher-order taxa (below) in the aquatic subsets.

At the level of major taxonomic units, all four sample groups were dominated by the same phylogenetic groups, both in prokaryotes and microeukaryotes (Figure 3). Gammaproteobacteria, Bacteroidia and Alphaproteobacteria were the most abundant classified prokaryotes, making up 26-31% of the ASV abundances found in the aquatic communities and 53-76% in the *A. anser* samples. However, some taxa such as Bacilli and Thermoleophilia were abundant in the aquatic communities in both years (9-10% together), but were either completely missing (Bacilli in 2018) or represented only with very low abundances (0.03-0.3% together) in the *A. anser* droppings. In contrast, the relative abundance of Gammaproteobacteria was higher in the *A. anser* droppings (41-60%) compared to the aquatic community samples (7-8%, Figure 3).

In microeukaryotes, Chlorophyta, Apicomplexa and Fungi were the most abundant among the classified taxonomic groups, altogether representing 29-31% of the ASV abundances in the aquatic communities and 20-29% in the *A. anser* droppings. There were also several groups that were abundant in the aquatic communities in both years (15-22% together) but were not characteristic in the *A. anser* samples (0-0.3% together), e.g. Cercozoa, Katablepharidophyta and Ochrophyta. However, Opalozoa was represented with higher abundance in *A. anser* samples (9-19%) than in the aquatic communities (0.2-1%) (Figure 3).

On the PCoA plots, the separation of prokaryotic samples according to sample type (aquatic communities against *A. anser* droppings) was more explicit compared to the microeukaryotic samples (Figure 4). At the same time, the PERMANOVA tests resulted in a significant effect of sample type in both cases (Table 1). Both prokaryotic and microeukaryotic samples were less separated by year (Figure 4), which was in line with the stronger effect (indicated by higher R^2^ values) of sample type compared to year (though both were significant) based on PERMANOVA tests (Table 1). Pairwise comparisons of the four sample groups showed similar significant differences with overall higher R^2^ values for pairs of different sample types in prokaryotes, while in microeukaryotes the difference was significant only for the pairs of different sample types (*A. anser* or aquatic communities; Table 1). A subsequent SIMPER analysis (Table S12) showed that the ASVs most responsible for these differences belonged to the dominant higher order taxa (Figure 3) and there was a complete overlap between the ASVs most responsible for the differences in sample type and sampling year (Table S12). The general patterns in the PCoA, PERMANOVA and pairwise comparisons repeated for the incidence and unselected data subsets were highly similar in both prokaryotes and microeukaryotes with clearer differences among sample types and sampling years (Table S13–S14, Figure S3–S5).

**Table 1.** Results of PERMANOVA and pairwise comparison performed on the aquatic subset (relative abundance data, Bray-Curtis dissimilarity; permutations=2000) of prokaryotic and microeukaryotic communities detected in aquatic community (AC) and *Anser anser* dropping samples in 2017 and 2018.

| Prokaryotes |  |  |  |  |  |  |  | Microeukaryotes |  |  |  |  |  |  |
| --- | --- | --- | --- | --- | --- | --- | --- | --- | --- | --- | --- | --- | --- | --- |
| Factors | Df | SS | MS | F | R<sup>2</sup> | p |  | Df | SS | MS | F | R<sup>2</sup> | p |  |
| Sample type | 1 | 3.977 | 3.977 | 11.548 | 0.125 | 0.0005 | *** | 1 | 0.929 | 0.929 | 2.058 | 0.056 | 0.0005 | *** |
| Sampling year | 1 | 1.684 | 1.685 | 4.892 | 0.053 | 0.0005 | *** | 1 | 0.592 | 0.592 | 1.312 | 0.036 | 0.0295 | * |
| Sample type*sampling year | 1 | 1.274 | 1.274 | 3.699 | 0.040 | 0.0015 | ** | 1 | 0.511 | 0.511 | 1.133 | 0.031 | 0.1894 | ns |
| Residuals | 72 | 24.794 | 0.344 |  | 0.781 |  |  | 32 | 14.441 | 0.451 |  | 0.877 |  |  |
| Total | 75 | 31.729 |  |  | 1.000 |  |  | 35 | 16.473 |  |  | 1.000 |  |  |

| Pairs | Df | SS | F | R<sup>2</sup> | p | P<sub>adj</sub> |  | Df | SS | F | R<sup>2</sup> | p | P<sub>adj</sub> |  |
| --- | --- | --- | --- | --- | --- | --- | --- | --- | --- | --- | --- | --- | --- | --- |
| 2017 *A. anser* - 2017 AC | 1 | 2.805 | 8.325 | 0.188 | 0.0005 | 0.0030 | ** | 1 | 0.734 | 1.672 | 0.095 | 0.0020 | 0.0120 | * |
| 2017 *A. anser* - 2018 *A. anser* | 1 | 1.754 | 4.701 | 0.116 | 0.0005 | 0.0030 | ** | 1 | 0.624 | 1.355 | 0.078 | 0.0745 | 0.4468 | ns |
| 2017 *A. anser* - 2018 AC | 1 | 3.486 | 11.512 | 0.242 | 0.0005 | 0.0030 | ** | 1 | 0.746 | 1.673 | 0.095 | 0.0025 | 0.0150 | * |
| 2017 AC - 2018 *A. anser* | 1 | 2.175 | 5.636 | 0.135 | 0.0005 | 0.0030 | ** | 1 | 0.775 | 1.697 | 0.096 | 0.0005 | 0.0030 | ** |
| 2017 AC - 2018 AC | 1 | 1.204 | 3.815 | 0.096 | 0.0005 | 0.0030 | ** | 1 | 0.480 | 1.085 | 0.063 | 0.2824 | 1.0000 | ns |
| 2018 *A. anser* - 2018 AC | 1 | 2.445 | 6.951 | 0.162 | 0.0005 | 0.0030 | ** | 1 | 0.706 | 1.522 | 0.087 | 0.0005 | 0.0030 | ** |

**Figure 4.**
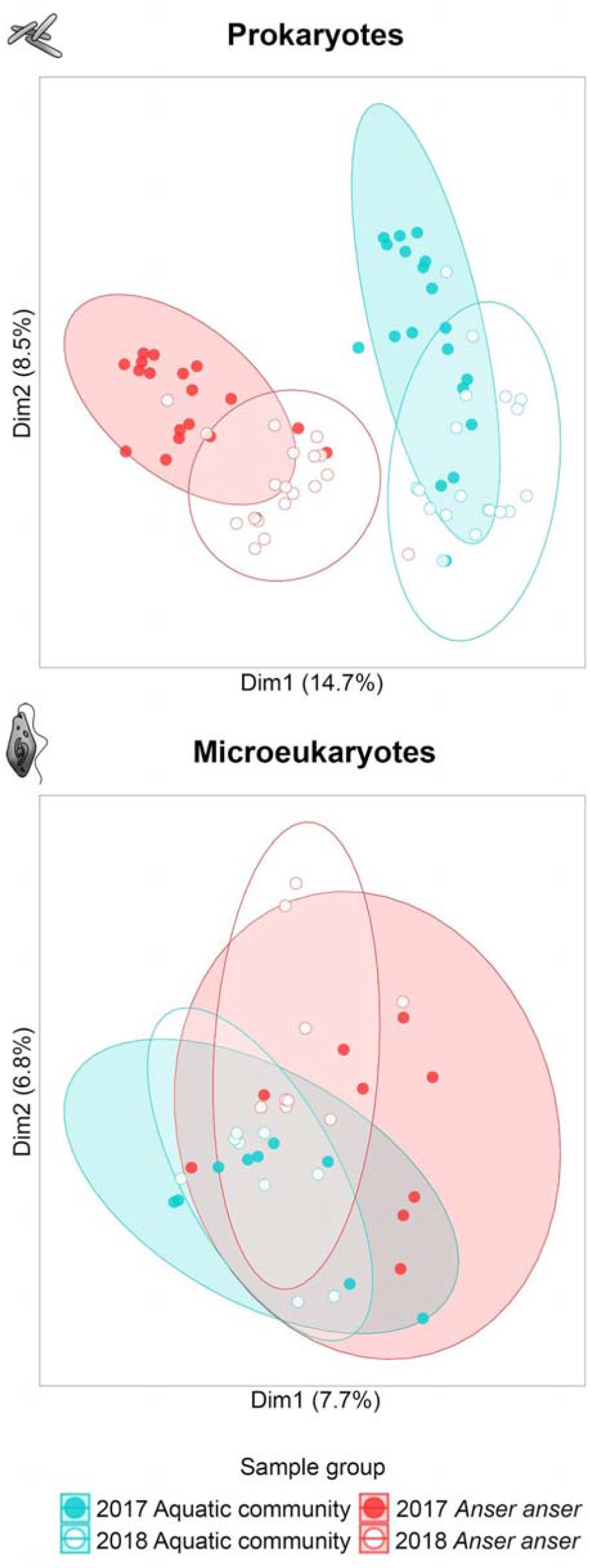
PCoA biplot of aquatic community and *Anser anser* dropping samples collected in 2017 and 2018. The analysis is based on the aquatic subset (relative abundance data, Bray-Curtis dissimilarity) of prokaryotic and microeukaryotic communities.

We finally compared the richness (Figure 5, Figure S6) and composition (Figure S7– S10) of microbes detected in the droppings of four waterbird species. Similar to the results based on rarefaction for *A. anser* droppings (Figure 2–3), only a fraction of the total species pool was recaptured in each waterbird species, but the actual proportion changed with species. The shorebirds, *C. pugnax* transported a similar fraction of microeukaryotic ASVs as geese (*A. anser*), both as individuals (mean richness) and collectively (the latter evidenced by regional extrapolated richness). Compared to them, *S. clypeata* proved to be much more efficient dispersal agents for microeukaryotes, dispersing almost twice as many ASVs as a same-sized group of any of the other species (Figure 5). Furthermore, we essentially found a similar number of microeukaryotic ASVs per *S. clypeata* dropping as in a random aquatic sample (Figure 5). PCoA ordinations both with abundance- and incidence-based data also showed a clear separation of *S. clypeata* from the rest of the waterbirds (Figure S7–S8).

**Figure 5.**
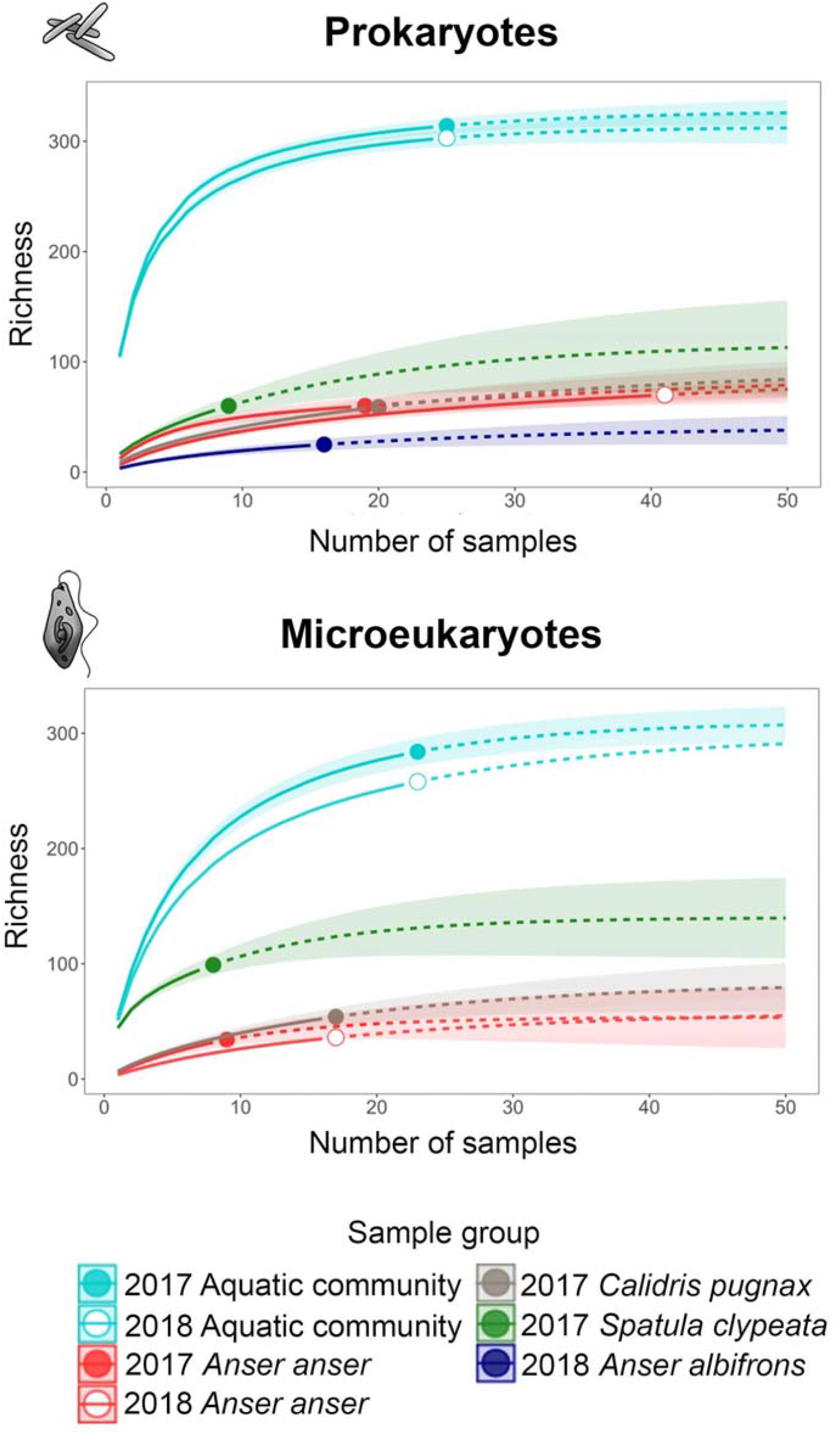
Accumulation curves with extrapolated ASV richness estimates (dashed lines) and 95% confidence intervals for the aquatic subsets of prokaryotes and microeukaryotes detected in the droppings of four waterbird species compared to the aquatic communities.

The comparison of waterbird species yielded somewhat different results for prokaryotes, where *S. clypeata* droppings no longer hosted significantly higher ASV richness than most of the other species (except for *A. albifrons*), and showed a large compositional overlap with communities potentially dispersed by shorebirds (*C. pugnax*) (Figure S7). Due to low read numbers for microeukaryotes in case of *A. albifrons*, in the two goose species, *A. anser* and *A. albifrons*, we could only compare the composition of prokaryotes in their droppings, where the difference we found was negligible (Figure S7-S8). While the overall composition of the detected set of prokaryotes was very similar among individual birds (Figure S7–S8), *A. albifrons* collectively transported a significantly lower diversity of ASVs: approximately only the half of those found in *A. anser* droppings (Figure 5).

In the unselected datasets, the prokaryotic and microeukaryotic communities transported by different bird species were much more distinct (Figure S9–S10), but even there, *A. anser* and *A. albifrons* samples showed high similarity.

## Discussion

The main novelty of our study is twofold. First, it represents the first comprehensive study on the role waterbirds play in the dispersal of aquatic microorganisms using DNA metabarcoding targeting communities of prokaryotes and unicellular microeukaryotes. We provided evidence that waterbirds potentially can disperse all major aquatic groups from bacteria through phytoplankton to protozoa. Second, we directly compared microorganisms that may be dispersed by waterbirds to the source pool (natural aquatic communities), thereby being able to investigate the share and identity of aquatic microbes readily transported by waterbirds. In this confined set of aquatic habitats (i.e., metacommunity), we indeed found a considerable share of aquatic communities detected in waterbird droppings. Although the difference among sample types (aquatic communities and *A. anser* droppings) was in general more conspicuous, the actual set of prokaryote ASVs detected in the bird droppings also showed differences between the two years, where the potentially dispersed set of microbes reflected the actual aquatic communities. This provided further evidence for the dispersal potential of waterbirds. Finally, the communities detected in the droppings of different waterbird species showed high similarities (regardless of their lifestyle), with a number of specific differences. The implications of our results showed minor sensitivity to the selection methods (unselected dataset or aquatic subset) or data type (abundance or incidence), and were largely consistent across prokaryotes and microeukaryotes. Altogether, our study provided the first explicit quantitative evidence clearly supporting that waterbirds are so far overlooked, yet potentially important dispersal agents of natural communities of aquatic microorganisms.

Prokaryotic and microeukaryotic communities of the aquatic subset were typical for soda lakes and pans of the region (Sinclair et al., 2015; Szabó et al., 2017, 2020). We found that 28% of the prokaryotic and 19% of the aquatic microeukaryotic ASVs were also present in the droppings of the dominant waterbirds species of the region, *A. anser*. Instead of dispersing a single or only a limited number of aquatic taxa, most of the major taxonomic groups of the aquatic communities were well-represented in the *A. anser* droppings. In waterbirds, gut retention time is short (Brochet, Guillemain, Gauthier-Clerc et al., 2010; Sánchez et al., 2012), which can contribute to a large share of undigested microorganisms. In extreme cases, even live plants (Silva et al., 2018), diatoms (Atkinson, 1971, 1980), aquatic invertebrates (Frisch et al., 2007; Green & Sánchez, 2006) and gelatinous fish eggs (Lovas-Kiss et al., 2020) can survive waterbird gut passage. Compared to them, the survival of microorganisms should be even higher, given their evolutionary adaptations to adverse conditions such as extreme values of pH, desiccation or UV radiation (Potts, 1999; Rainey et al., 2005; Schleper et al., 1995). Even though we did not test the viability of the detected microbes directly, these altogether make it highly likely that the ASVs we found included viable cells and hence indicate the possibility of successful dispersal events.

We found that community composition of microbes, i.e., both the prokaryotic and microeukaryotic communities in the aquatic samples, and the dispersed ASV set in *A. anser* droppings were different between the two years. Besides, the difference in aquatic prokaryotic communities was also reflected by the communities found in the droppings (as evidenced by the PCoA plots). Theses altogether indicate that the set of potentially dispersed prokaryotes reflects the natural microbial communities available in the local aquatic habitats at the given time. That is, our observations confirm the previous assumption that internal dispersal depends on the availability of aquatic (food) organisms (e.g. Brochet, Guillemain, Fritz, et al., 2010; Frisch et al., 2007), which can vary in time and is facilitated by the weak digestion efficiency mentioned above.

Even though *A. anser* do not feed directly from the water but rather consume seeds, stems and leaves of aquatic macrophytes and terrestrial plants (Middleton & van der Valk, 1987), they can pick up microbes while drinking and while feeding on aquatic macrophytes or even while preening their damp feathers after bathing. Our results showed that this feeding mode still makes them potentially efficient dispersal agents for aquatic microbial metacommunities. At the same time, we showed a high heterogeneity of prokaryotic and microeukaryotic ASV composition across bird droppings, indicating stochastic pickup by the individual birds. Although the difference in local and regional richness between droppings and aquatic communities was still remarkable, the compositional variation among droppings was moderated when *A. anser* gut microbiota was also considered, leading us to the conclusion that the gut microorganism composition of *A. anser* is specific to the species. This is in line with the findings of Laviad-Shitrit et al. (2019) that waterbird species host unique gut bacterial communities.

We found that not only *A. anser* but the other three bird species can also transport a considerable share of the natural microbial communities present in the ponds. While we found some differences between the waterbird species, these were not completely congruent with their feeding habits and habitat use. In spite of the terrestrial feeding habit of *A. anse*r, the number of ASVs transported by them was largely comparable to those found in shorebirds, *C. pugnax*, which prefer to feed in the shallow shoreline regions of ponds (Baccetti et al., 1998) and may directly consume biofilm communities as shown for multiple *Calidris* spp. (Kuwae et al., 2008, 2012). According to our results, they all can disperse quite similar microeukaryotic communities across aquatic habitats.

When considering the potentially dispersed prokaryotes, we did not find remarkable differences among the different bird species, neither in ASV richness, nor in composition. *A. anse*r and *A. albifrons* transported quite similar prokaryotic communities and their gut microbiome also seems to be largely the same, which is not surprising given that both have a predominantly terrestrial herbivorous feeding habit (Ely & Raveling, 2011; Middleton & van der Valk, 1987). Nevertheless, of the two, *A. anser* hosted a higher number of ASVs in their droppings, which implies that it might have a more important role in the endozoochory of prokaryotes.

The only species that showed marked differences from the rest of the waterbirds was *S. clypeata*. They not only transported different microeukaryotic communities but also captured a much larger fraction of the aquatic source pool, therefore they can be considered as the most effective dispersal agents. However, in terms of transporting prokaryotes, they were no longer so prominent. A reasonable explanation for our observations can be that *S. clypeata*, unlike the other waterbirds we studied, is a planktivore species sieving plankton from the open water (Matsubara et al., 1994). The low interlamellar distances in its specialized spoon-shaped bill enable an effective accumulation of aquatic microorganisms even smaller than 500 µm (Gurd, 2007; Kooloos et al., 1989). Thus, microeukaryotes and their propagules of this size can be easily captured and concentrated, whereas bacterioplankton with a smaller size fraction probably flows through their lamellae.

Altogether, our results are based on a representative comparison of equal sample sizes across aquatic habitats and bird droppings. We proved that within small-scale pond and lake networks (10-20 km), waterbirds can be important dispersal agents of both prokaryotes and microeukaryotes, given that the spatial scale of such pondscapes coincides with the local daily movements of waterbirds, including *A. anser* (Bell, 1988; Boos et al., 2019; Link et al., 2011; Nilsson & Persson, 1992). As the study region might host up to hundreds of thousands of waterbirds (Dick et al., 1994), which themselves might defecate even up to 80 times per day (Oláh, 2003; Sterbetz, 1992), their overall contribution to biotic connectivity is expected to be immense, eventually being able to transport most members of the aquatic microbial metacommunity among the habitats.

Finally, this study also has important implications for larger spatial scales. According to their flight speed and gut retention times, waterbirds are able to transport their intestinal contents over thousands of kilometers during their migration (Viana et al., 2013a, 2013b, 2016), and therefore can be important dispersal agents of aquatic microorganisms not only on regional but even on continental scales. By dispersing microorganisms, they can have a significant role in forming biodiversity patterns and sustaining ecosystem functions where the importance of microbes is indisputable (Bell et al., 2005; Graham et al., 2016; Wohl et al., 2004).

## Acknowledgements

Authors thank the colleagues of Lake Neusiedl Biological Station, Thomas Zechmeister, Richard Haider and Rudolf Schalli, for their help in field work and providing access to their labs; colleagues of Fertő-Hanság and Neusiedler See - Seewinkel National Parks for providing and helping us access the sites as part of the Vogelwarte - Madárvárta 2 project; Christian Preiler for further practical help; Mia Bengtsson for her help with selecting the primers used in the study; Lucie Zinger and two anonymous reviewers whose comments helped to improve the manuscript. The study was supported by the Interreg V□A Austria□Hungary programme of the European Regional Development Fund (Vogelwarte Madárvárta 2). Field work was permitted through the Fertő-Hanság and Neusiedler See - Seewinkel National Parks as part of the project and additionally through the permits ND-10-05-4326-4-2017 (Neusiedl am See Bezirkshauptmannschaft) and PE-KTF/882-2/2018 (Pest Megyei Kormányhivatal Környezetvédelmi és Természetvédelmi Főosztály). BS acknowledges further support by NKFIH-132095. ZH and CFV were supported by GINOP 2.3.2.□15□2016□00057 and the AQUACOSM-plus project of the European Union’s Horizon 2020 research and innovation programme under grant agreement No. 871081. ZH acknowledges support from the Janos Bolyai Research Scholarship of the Hungarian Academy of Sciences. AS was supported by the Wenner-Gren Foundations. BS, AS, CFV, ZM, and ZH were furthermore supported by the NKFIH-471-3/2021 project.

## Data availability statement

Raw sequence reads were deposited in the NCBI SRA database and are accessible through the BioProject accession PRJNA748202.

## Biosketch

Beáta Szabó is a postdoc researcher at Institute of Aquatic Ecology, Centre for Ecological Research and interested in the processes underlying the formation of biodiversity and metacommunities of aquatic microorganisms with a special focus on small temporary ponds.

## Author contributions

ZH, CFV and RP conceived the ideas; ZH, CFV, EB and DL collected the samples; ZM performed the lab work; AS and ZM analysed the molecular data; BS and ZH performed the statistical analyses; BS wrote the manuscript with significant contributions by ZH, AS and CFV; all authors commented on earlier versions of the manuscript.

## Supporting information

Additional supporting information may be found in the online version of the article at the publisher’s website.

