## Supporting information for "Microbial stowaways – waterbirds as dispersal vectors of aquatic pro- and microeukaryotic communities"

This file includes the following tables and figures:

Table S1, Tables S10-S14

Figures S1-S10

The following other supplementary materials for this manuscript are available as separate Excel spreadsheets:

Tables S2-S9

Table S1. Number of samples after re-sampling the amplicon data per sample group, together with the feeding mode (Roland 2019) of the studied waterbird species.

|  |  |  | **Prokaryotes** | | **Microeukaryotes** | |
| --- | --- | --- | --- | --- | --- | --- |
|  |  | **Feeding habit** | Unselected dataset | Aquatic subset | Unselected dataset | Aquatic subset |
| 2017 | *A. anser* | terrestrial grazing | 26 | 19 | 10 | 9 |
|  | *C. pugnax* | mud-probing in the shallow shoreline regions | 20 | 20 | 17 | 17 |
|  | *R. avosetta* | scything in the shallow shoreline regions | 4 | 4 | 3 | 3 |
|  | *S. clypeata* | filter-feeding in the open water | 9 | 9 | 8 | 8 |
|  | Aquatic community | - | 25 | 25 | 23 | 23 |
| 2018 | *A. albifrons* | terrestrial grazing | 18 | 16 | - | - |
|  | *A. anser* | terrestrial grazing | 44 | 41 | 25 | 17 |
|  | Aquatic community | - | 25 | 25 | 23 | 23 |

Table S2. 16S ASV set rarefied to 8620 read per sample (referred to as unselected prokaryotic dataset). Sampling year and sample group (five waterbird species and aquatic community) are indicated. (Excel file available as a separate document).

Table S3. 18S ASV set rarefied to 2432 read per sample (referred to as unselected microeukaryotic dataset). Sampling year and sample group (five waterbird species and aquatic community) are indicated. (Excel file available as a separate document).

Table S4. Set of 16S ASVs present at least in one aquatic community sample with ≥1% relative abundance (referred to as aquatic subset of prokaryotes). Sampling year and sample group (five waterbird species and aquatic community) are indicated. (Excel file available as a separate document).

Table S5. Set of 18S ASVs present at least in one aquatic community sample with ≥1% relative abundance (referred to as aquatic subset of microeukaryotes). Sampling year and sample group (five waterbird species and aquatic community) are indicated. (Excel file available as a separate document).

Table S6. List of taxa assigned to the unselected 16S ASV set. (Excel file available as a separate document).

Table S7. List of taxa assigned to the unselected 18S ASV set. (Excel file available as a separate document).

Table S8. List of taxa assigned to the aquatic subset of 16S ASVs. (Excel file available as a separate document).

Table S9. List of taxa assigned to the aquatic subset of 18S ASVs. (Excel file available as a separate document).

Table S10. Results of Scheirer–Ray–Hare test and Dunn’s post-hoc test for pairwise comparisons performed on the α- and β-diversity of the prokaryotic and microeukaryotic aquatic subsets in aquatic communities (AC) and *Anser anser* droppings in 2017 and 2018.

|  |  | **Prokaryotes** | | | | | **Microeukaryotes** | | | | |
| --- | --- | --- | --- | --- | --- | --- | --- | --- | --- | --- | --- |
|  | Factors | Df | SS | H | p |  | Df | SS | H | p |  |
| α-diversity | Sample type | 1 | 27436.0 | 56.323 | 0.0000 | ******* | 1 | 2916.0 | 26.389 | 0.0000 | ******* |
|  | Sampling year | 1 | 90.6 | 0.186 | 0.6662 | ns | 1 | 16.0 | 0.145 | 0.7036 | ns |
|  | Sample type*sampling year | 1 | 82.1 | 0.169 | 0.6814 | ns | 1 | 9.0 | 0.081 | 0.7753 | ns |
|  | Residuals | 72 | 8925.2 |  |  |  | 32 | 926.5 |  |  |  |
| β-diversity | Sample type | 1 | 12662.6 | 25.976 | 0.0000 | ******* | 1 | 1111.11 | 10.024 | 0.0015 | ****** |
|  | Sampling year | 1 | 521.1 | 1.069 | 0.3012 | ns | 1 | 11.11 | 0.100 | 0.7515 | ns |
|  | Sample type*sampling year | 1 | 796.3 | 1.633 | 0.2012 | ns | 1 | 49.00 | 0.442 | 0.5061 | ns |
|  | Residuals | 72 | 22580.5 |  |  |  | 32 | 2708.28 |  |  |  |
|  | Pairs | Z | p | p_adj_ |  |  | Z | p | p_adj_ |  |  |
| α-diversity | 2017 *A. anser* - 2017 AC | -5.016 | 5.265e-07 | 1.053e-06 | ******* |  | -3.431 | 0.0006 | 0.0012 | ****** |  |
|  | 2017 *A. anser* - 2018 *A. anser* | 0.595 | 0.5516 | 0.6619 | ns |  | 0.471 | 0.6377 | 0.7653 | ns |  |
|  | 2017 *A. anser* - 2018 AC | -5.002 | 5.682e-07 | 8.524e-07 | ******* |  | -3.363 | 0.0008 | 0.0012 | ****** |  |
|  | 2017 AC - 2018 *A. anser* | 5.612 | 2.003e-08 | 1.202e-07 | ******* |  | 3.902 | 0.0001 | 0.0006 | ******* |  |
|  | 2017 AC - 2018 AC | 0.015 | 0.9883 | 0.9883 | ns |  | 0.067 | 0.9464 | 0.9464 | ns |  |
|  | 2018 *A. anser* - 2018 AC | -5.597 | 2.180e-08 | 6.540e-08 | ******* |  | -3.834 | 0.0001 | 0.0004 | ******* |  |
| β-diversity | 2017 *A. anser* - 2017 AC | 2.700 | 0.0069 | 0.0104 | ***** |  | 1.769 | 0.0770 | 0.1154 | ns |  |
|  | 2017 *A. anser* - 2018 *A. anser* | -1.635 | 0.1021 | 0.1225 | ns |  | -0.694 | 0.4877 | 0.5852 | ns |  |
|  | 2017 *A. anser* - 2018 AC | 2.873 | 0.0041 | 0.0081 | ****** |  | 2.015 | 0.0439 | 0.0878 | **.** |  |
|  | 2017 AC - 2018 *A. anser* | -4.335 | 1.458e-05 | 4.374e-05 | ******* |  | -2.463 | 0.0138 | 0.0414 | ***** |  |
|  | 2017 AC - 2018 AC | 0.173 | 0.8629 | 0.8629 | ns |  | 0.246 | 0.8055 | 0.8055 | ns |  |
|  | 2018 *A. anser* - 2018 AC | 4.508 | 6.556e-06 | 3.934e-05 | ******* |  | 2.709 | 0.0068 | 0.0405 | ***** |  |

Table S11. Results of Scheirer–Ray–Hare test and Dunn’s post-hoc test for pairwise comparisons performed on the α- and β-diversity of the unselected prokaryotic and microeukaryotic communities in aquatic communities (AC) and *Anser anser* droppings in 2017 and 2018.

|  |  | **Prokaryotes** | | | | | **Microeukaryotes** | | | | |
| --- | --- | --- | --- | --- | --- | --- | --- | --- | --- | --- | --- |
|  | Factors | Df | SS | H | p |  | Df | SS | H | p |  |
| α-diversity | Sample type | 1 | 32364 | 38.453 | 0.0000 | *** | 1 | 3027.6 | 22.164 | 0.0000 | *** |
|  | Sampling year | 1 | 1218 | 1.447 | 0.2290 | ns | 1 | 102.4 | 0.750 | 0.3866 | ns |
|  | Sample type*sampling year | 1 | 2852 | 3.388 | 0.0657 | . | 1 | 96.1 | 0.704 | 0.4016 | ns |
|  | Residuals | 96 | 46890 |  |  |  | 36 | 2101.4 |  |  |  |
| β-diversity | Sample type | 1 | 6209 | 7.378 | 0.0066 | ** | 1 | 240.1 | 1.757 | 0.1849 | ns |
|  | Sampling year | 1 | 502 | 0.596 | 0.4401 | ns | 1 | 60.0 | 0.439 | 0.5074 | ns |
|  | Sample type*sampling year | 1 | 29 | 0.035 | 0.8523 | ns | 1 | 18.2 | 0.133 | 0.7149 | ns |
|  | Residuals | 96 | 76585 |  |  |  | 36 | 5009.7 |  |  |  |
|  | Pairs | Z | p | p_adj_ |  |  | Z | p | p_adj_ |  |  |
| α-diversity | 2017 *A. anser* - 2017 AC | -5.686 | 1.298e-08 | 7.788e-08 | *** |  | -3.922 | 8.781e-05 | 0.0003 | *** |  |
|  | 2017 *A. anser* - 2018 *A. anser* | -0.451 | 0.6521 | 0.6521 | ns |  | 0.019 | 0.9847 | 0.9847 | ns |  |
|  | 2017 *A. anser* - 2018 AC | -3.534 | 0.0004 | 0.0008 | *** |  | -2.717 | 0.0066 | 0.0099 | ** |  |
|  | 2017 AC - 2018 *A. anser* | 5.235 | 1.646e-07 | 4.938e-07 | *** |  | 3.941 | 8.109e-05 | 0.0004 | *** |  |
|  | 2017 AC - 2018 AC | 2.152 | 0.0314 | 0.0377 | * |  | 1.2053 | 0.2281 | 0.2737 | ns |  |
|  | 2018 *A. anser* - 2018 AC | -3.083 | 0.0020 | 0.0031 | ** |  | -2.736 | 0.0062 | 0.0124 | * |  |
| β-diversity | 2017 *A. anser* - 2017 AC | 2.052 | 0.0401 | 0.1204 | ns |  | 1.196 | 0.2318 | 0.6955 | ns |  |
|  | 2017 *A. anser* - 2018 *A. anser* | -0.414 | 0.6786 | 0.6786 | ns |  | 0.727 | 0.4672 | 0.9345 | ns |  |
|  | 2017 *A. anser* - 2018 AC | 1.375 | 0.1692 | 0.2539 | ns |  | 1.406 | 0.1597 | 0.9581 | ns |  |
|  | 2017 AC - 2018 *A. anser* | -2.467 | 0.0136 | 0.0818 | . |  | -0.469 | 0.6393 | 0.7671 | ns |  |
|  | 2017 AC - 2018 AC | -0.678 | 0.4980 | 0.5976 | ns |  | 0.210 | 0.8333 | 0.8333 | ns |  |
|  | 2018 *A. anser* - 2018 AC | 1.789 | 0.0736 | 0.1472 | ns |  | 0.679 | 0.4970 | 0.7456 | ns |  |

Table S12. 15 ASVs most responsible for the dissimilarities among sample types and sampling years based on SIMPER analysis of the aquatic subset (abundance data, Bray-Curtis dissimilarity).

|  | **Prokaryotes** | | | | **Microeukaryotes** | | | |
| --- | --- | --- | --- | --- | --- | --- | --- | --- |
|  | ASV | Taxon | Taxlevel 3 | Cum. contr. (%) | ASV | Taxon | Taxlevel 3 | Cum. contr. (%) |
| Sample type | ASV31427 | Idiomarinaceae_unclass | Gammaproteobacteria | 8.73 | ASV08172 | Eimeriidae_unclass | Apicomplexa | 5.25 |
|  | ASV25904 | Comamonadaceae_unclass | Gammaproteobacteria | 13.90 | ASV04489 | Opalozoa_unclass | Opalozoa | 10.37 |
|  | ASV19092 | WD2101_soil_group_unclass | Phycisphaerae | 17.79 | ASV01901 | Spermatozopsis_exsultans | Chlorophyta | 14.88 |
|  | ASV14049 | Anaerolineaceae_unclass | Anaerolineae | 21.05 | ASV05136 | Eukaryota_unclass | Eukaryota_unclass | 19.32 |
|  | ASV12701 | Burkholderiales_unclass | Gammaproteobacteria | 23.63 | ASV09777 | Eukaryota_unclass | Eukaryota_unclass | 22.97 |
|  | ASV30974 | Devosiaceae_unclass | Alphaproteobacteria | 26.00 | ASV05057 | Eukaryota_unclass | Eukaryota_unclass | 26.40 |
|  | ASV06229 | Ardenticatenales_unclass | Anaerolineae | 28.18 | ASV03275 | Planctonema_sp. | Chlorophyta | 29.45 |
|  | ASV01228 | PLTA13_unclass | Gammaproteobacteria | 30.22 | ASV06048 | Eukaryota_unclass | Eukaryota_unclass | 32.49 |
|  | ASV22608 | Gaiella_unclass | Thermoleophilia | 32.20 | ASV02445 | Eimeriida_unclass | Apicomplexa | 35.43 |
|  | ASV10140 | Bacteria_unclass | Bacteria_unclass | 34.14 | ASV01317 | Eukaryota_unclass | Eukaryota_unclass | 38.25 |
|  | ASV06432 | Geminicoccaceae_unclass | Alphaproteobacteria | 35.81 | ASV03490 | Blastocystis_ST7 | Opalozoa | 40.55 |
|  | ASV12089 | Beijerinckiaceae_unclass | Alphaproteobacteria | 37.38 | ASV07062 | Eimeriida_unclass | Apicomplexa | 42.54 |
|  | ASV13101 | Cytophagales_unclass | Bacteroidia | 38.89 | ASV09281 | Eukaryota_unclass | Eukaryota_unclass | 44.40 |
|  | ASV14582 | Microbacteriaceae_unclass | Actinobacteria | 40.39 | ASV08226 | Eukaryota_unclass | Eukaryota_unclass | 46.08 |
|  | ASV02360 | Bacteriovoracaceae_unclass | Bdellovibrionia | 41.80 | ASV04491 | Pyramimonas_gelidicola | Chlorophyta | 47.76 |
| Sampling year | ASV31427 | Idiomarinaceae_unclass | Gammaproteobacteria | 8.63 | ASV08172 | Eimeriidae_unclass | Apicomplexa | 5.53 |
|  | ASV25904 | Comamonadaceae_unclass | Gammaproteobacteria | 13.55 | ASV04489 | Opalozoa_unclass | Opalozoa | 10.72 |
|  | ASV19092 | WD2101_soil_group_unclass | Phycisphaerae | 17.46 | ASV05136 | Eukaryota_unclass | Eukaryota_unclass | 15.04 |
|  | ASV14049 | Anaerolineaceae_unclass | Anaerolineae | 20.81 | ASV01901 | Spermatozopsis_exsultans | Chlorophyta | 19.28 |
|  | ASV12701 | Burkholderiales_unclass | Gammaproteobacteria | 23.50 | ASV09777 | Eukaryota_unclass | Eukaryota_unclass | 22.96 |
|  | ASV30974 | Devosiaceae_unclass | Alphaproteobacteria | 25.92 | ASV05057 | Eukaryota_unclass | Eukaryota_unclass | 26.47 |
|  | ASV06229 | Ardenticatenales_unclass | Anaerolineae | 28.19 | ASV06048 | Eukaryota_unclass | Eukaryota_unclass | 29.60 |
|  | ASV01228 | PLTA13_unclass | Gammaproteobacteria | 30.26 | ASV03275 | Planctonema_sp. | Chlorophyta | 32.62 |
|  | ASV22608 | Gaiella_unclass | Thermoleophilia | 32.29 | ASV02445 | Eimeriida_unclass | Apicomplexa | 35.61 |
|  | ASV10140 | Bacteria_unclass | Bacteria_unclass | 34.14 | ASV01317 | Eukaryota_unclass | Eukaryota_unclass | 38.40 |
|  | ASV12089 | Beijerinckiaceae_unclass | Alphaproteobacteria | 35.80 | ASV03490 | Blastocystis_ST7 | Opalozoa | 40.57 |
|  | ASV13101 | Cytophagales_unclass | Bacteroidia | 37.39 | ASV07062 | Eimeriida_unclass | Apicomplexa | 42.59 |
|  | ASV14582 | Microbacteriaceae_unclass | Actinobacteria | 38.94 | ASV09281 | Eukaryota_unclass | Eukaryota_unclass | 44.40 |
|  | ASV06432 | Geminicoccaceae_unclass | Alphaproteobacteria | 40.46 | ASV08226 | Eukaryota_unclass | Eukaryota_unclass | 46.09 |
|  | ASV02360 | Bacteriovoracaceae_unclass | Bdellovibrionia | 41.89 | ASV04491 | Pyramimonas_gelidicola | Chlorophyta | 47.74 |

Table S13. Results of PERMANOVA and pairwise comparison performed on the incidence data subset (Sørensen dissimilarity; permutations=2000) of prokaryotic and microeukaryotic communities detected in aquatic community (AC) and *Anser anser* dropping samples in 2017 and 2018.

|  | **Prokaryotes** | | | | | | | **Microeukaryotes** | | | | | | |
| --- | --- | --- | --- | --- | --- | --- | --- | --- | --- | --- | --- | --- | --- | --- |
| Factors | Df | SS | MS | F | R^2^ | p |  | Df | SS | MS | F | R^2^ | p |  |
| Sample type | 1 | 6.172 | 6.172 | 25.715 | 0.239 | 0.0005 | ******* | 1 | 1.891 | 1.891 | 5.477 | 0.131 | 0.0005 | ******* |
| Sampling year | 1 | 1.407 | 1.407 | 5.862 | 0.054 | 0.0010 | ****** | 1 | 0.806 | 0.806 | 2.335 | 0.056 | 0.0010 | ****** |
| Sample type*sampling year | 1 | 1.012 | 1.012 | 4.215 | 0.039 | 0.0005 | ****** | 1 | 0.719 | 0.719 | 2.082 | 0.050 | 0.0010 | ****** |
| Residuals | 72 | 17.282 | 0.240 |  | 0.668 |  |  | 32 | 11.048 | 0.345 |  | 0.764 |  |  |
| Total | 75 | 25.873 |  |  | 1.000 |  |  | 35 | 14.464 |  |  | 1.000 |  |  |
| Pairs | Df | SS | F | R^2^ | p | p_adj_ |  | Df | SS | F | R^2^ | p | p_adj_ |  |
| 2017 *A. anser* - 2017 AC | 1 | 3.640 | 15.677 | 0.303 | 0.0005 | 0.0030 | ****** | 1 | 1.258 | 3.637 | 0.185 | 0.0005 | 0.0030 | ****** |
| 2017 *A. anser* - 2018 *A. anser* | 1 | 1.647 | 4.784 | 0.117 | 0.0005 | 0.0030 | ****** | 1 | 0.737 | 1.672 | 0.095 | 0.0060 | 0.0360 | ***** |
| 2017 *A. anser* - 2018 AC | 1 | 3.788 | 17.156 | 0.323 | 0.0005 | 0.0030 | ****** | 1 | 1.423 | 4.414 | 0.216 | 0.0005 | 0.0030 | ***** |
| 2017 AC - 2018 *A. anser* | 1 | 3.791 | 14.624 | 0.289 | 0.0005 | 0.0030 | ****** | 1 | 1.274 | 3.462 | 0.178 | 0.0005 | 0.0030 | ****** |
| 2017 AC - 2018 AC | 1 | 0.772 | 5.683 | 0.136 | 0.0010 | 0.0060 | ****** | 1 | 0.788 | 3.155 | 0.165 | 0.0005 | 0.0030 | ****** |
| 2018 *A. anser* - 2018 AC | 1 | 3.544 | 14.298 | 0.284 | 0.0005 | 0.0030 | ****** | 1 | 1.351 | 3.922 | 0.197 | 0.0005 | 0.0030 | ****** |

Table S14. Results of PERMANOVA and pairwise comparison performed on the unselected abundance and incidence dataset (Bray-Curtis and Sørensen dissimilarity, respectively; permutations=2000) of prokaryotic and microeukaryotic communities detected in aquatic community (AC) and *Anser anser* dropping samples in 2017 and 2018.

|  |  | **Prokaryotes** | | | | | | | **Microeukaryotes** | | | | | | |
| --- | --- | --- | --- | --- | --- | --- | --- | --- | --- | --- | --- | --- | --- | --- | --- |
| Bray-Curtis | Factors | Df | SS | MS | F | R^2^ | p |  | Df | SS | MS | F | R^2^ | p |  |
|  | Sample type | 1 | 6.050 | 6.050 | 16.447 | 0.139 | 0.0005 | ******* | 1 | 2.551 | 2.551 | 6.834 | 0.147 | 0.0005 | ******* |
|  | Sampling year | 1 | 1.056 | 1.056 | 2.872 | 0.024 | 0.0015 | ****** | 1 | 0.692 | 0.692 | 1.854 | 0.040 | 0.0165 | ***** |
|  | Sample type*sampling year | 1 | 0.965 | 0.965 | 2.623 | 0.022 | 0.0015 | ****** | 1 | 0.679 | 0.679 | 1.820 | 0.039 | 0.0275 | ***** |
|  | Residuals | 96 | 35.313 | 0.368 |  | 0.814 |  |  | 36 | 13.437 | 0.373 |  | 0.774 |  |  |
|  | Total | 99 | 43.384 |  |  | 1.000 |  |  | 39 | 17.359 |  |  | 1.000 |  |  |
|  | Pairs | Df | SS | F | R^2^ | p | p_adj_ |  | Df | SS | F | R^2^ | p | p_adj_ |  |
|  | 2017 *A. anser* - 2017 AC | 1 | 3.210 | 8.450 | 0.150 | 0.0005 | 0.0030 | ****** | 1 | 1.580 | 4.196 | 0.189 | 0.0005 | 0.0030 | ****** |
|  | 2017 *A. anser* - 2018 *A. anser* | 1 | 0.754 | 1.907 | 0.038 | 0.0055 | 0.0330 | ***** | 1 | 0.737 | 2.275 | 0.112 | 0.0155 | 0.0929 | **.** |
|  | 2017 *A. anser* - 2018 AC | 1 | 4.022 | 11.571 | 0.194 | 0.0005 | 0.0030 | ****** | 1 | 1.565 | 4.123 | 0.186 | 0.0005 | 0.0030 | ****** |
|  | 2017 AC - 2018 *A. anser* | 1 | 3.084 | 7.947 | 0.142 | 0.0005 | 0.0030 | ****** | 1 | 1.678 | 4.573 | 0.203 | 0.0005 | 0.0030 | ****** |
|  | 2017 AC - 2018 AC | 1 | 1.268 | 3.722 | 0.072 | 0.0005 | 0.0030 | ****** | 1 | 0.635 | 1.502 | 0.077 | 0.0125 | 0.0750 | **.** |
|  | 2018 *A. anser* - 2018 AC | 1 | 3.805 | 10.694 | 0.182 | 0.0005 | 0.0030 | ****** | 1 | 1.650 | 4.460 | 0.199 | 0.0005 | 0.0030 | ****** |
| Sørensen | Factors | Df | SS | MS | F | R^2^ | p |  | Df | SS | MS | F | R^2^ | p |  |
|  | Sample type | 1 | 6.328 | 6.328 | 18.770 | 0.155 | 0.0005 | ******* | 1 | 2.237 | 2.237 | 5.952 | 0.130 | 0.0005 | ******* |
|  | Sampling year | 1 | 1.203 | 1.203 | 3.569 | 0.030 | 0.0005 | ******* | 1 | 0.717 | 0.717 | 1.909 | 0.042 | 0.0035 | ***** |
|  | Sample type*sampling year | 1 | 0.892 | 0.892 | 2.645 | 0.022 | 0.0015 | ****** | 1 | 0.694 | 0.694 | 1.848 | 0.040 | 0.0165 | ***** |
|  | Residuals | 96 | 32.364 | 0.337 |  | 0.793 |  |  | 36 | 13.529 | 0.376 |  | 0.788 |  |  |
|  | Total | 99 | 40.787 |  |  | 1.000 |  |  | 39 | 17.178 |  |  | 1.000 |  |  |
|  | Pairs | Df | SS | F | R^2^ | p | p_adj_ |  | Df | SS | F | R^2^ | p | p_adj_ |  |
|  | 2017 *A. anser* - 2017 AC | 1 | 3.620 | 10.720 | 0.183 | 0.0005 | 0.0030 | ****** | 1 | 1.417 | 3.737 | 0.172 | 0.0005 | 0.0030 | ****** |
|  | 2017 *A. anser* - 2018 *A. anser* | 1 | 1.036 | 3.013 | 0.059 | 0.0005 | 0.0030 | ****** | 1 | 0.719 | 1.870 | 0.094 | 0.0005 | 0.0030 | ****** |
|  | 2017 *A. anser* - 2018 AC | 1 | 3.821 | 11.663 | 0.195 | 0.0005 | 0.0030 | ****** | 1 | 1.374 | 3.621 | 0.167 | 0.0005 | 0.0030 | ****** |
|  | 2017 AC - 2018 *A. anser* | 1 | 3.710 | 10.703 | 0.182 | 0.0005 | 0.0030 | ****** | 1 | 1.581 | 4.246 | 0.191 | 0.0005 | 0.0030 | ****** |
|  | 2017 AC - 2018 AC | 1 | 1.059 | 3.206 | 0.063 | 0.0005 | 0.0030 | ****** | 1 | 0.693 | 1.887 | 0.095 | 0.0010 | 0.0060 | ****** |
|  | 2018 *A. anser* - 2018 AC | 1 | 3.600 | 10.695 | 0.182 | 0.0005 | 0.0030 | ****** | 1 | 1.514 | 4.065 | 0.184 | 0.0005 | 0.0030 | ****** |

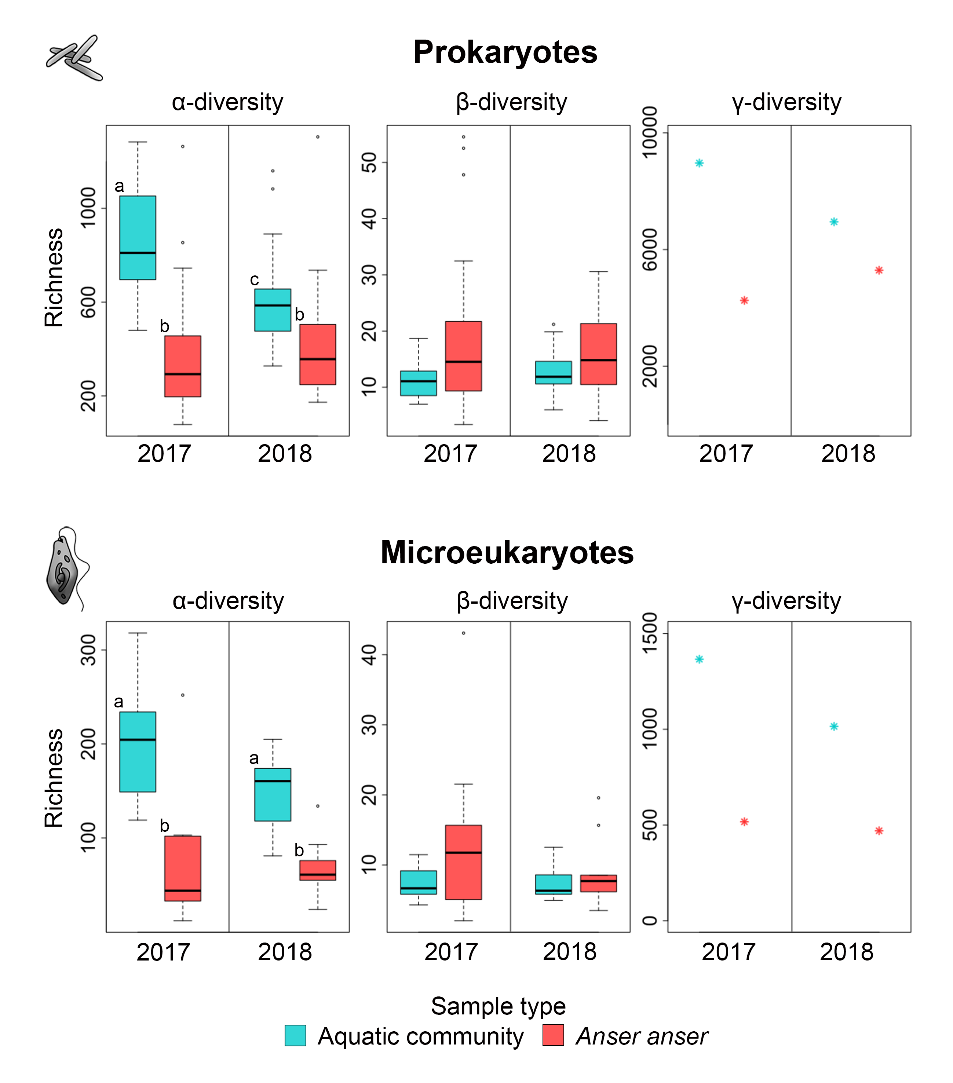

Figure S1. α-, β- and γ-diversity of the unselected prokaryotic and microeukaryotic communities in aquatic communities and *Anser anser* droppings in 2017 and 2018. Different letters indicate statistically significant differences at a significant level of p_adj_<0.05 based on Dunn’s pairwise post-hoc test. Pairwise gamma diversity comparisons are presented in Figure S6.

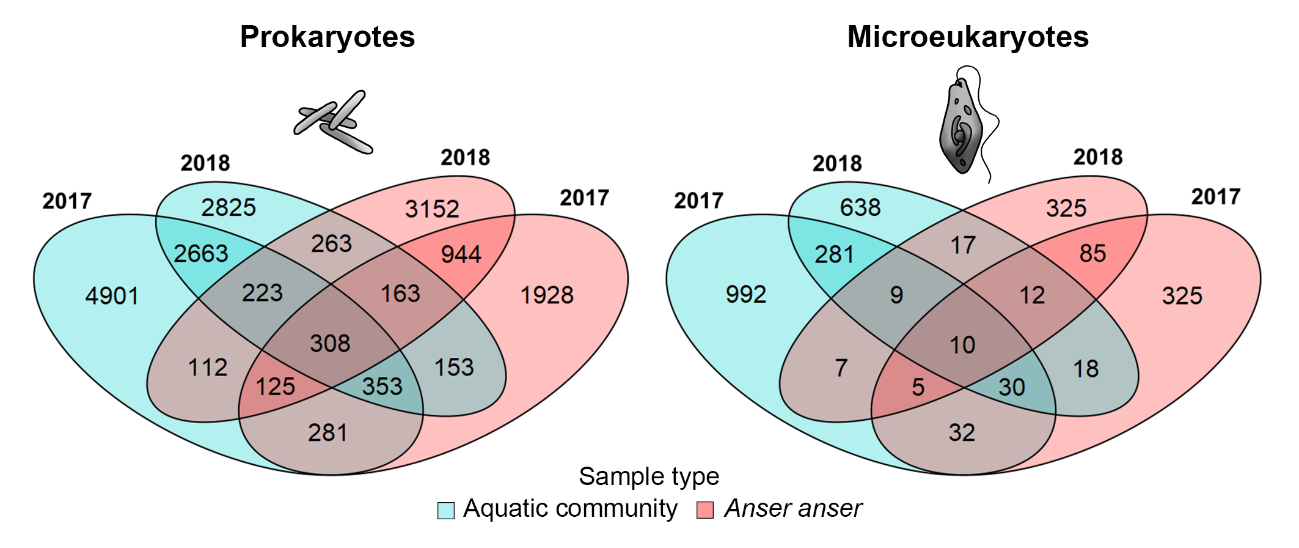

Figure S2. Number of prokaryotic and microeukaryotic ASVs shared among sample types (aquatic community and *Anser anser* dropping) and years in the unselected datasets.

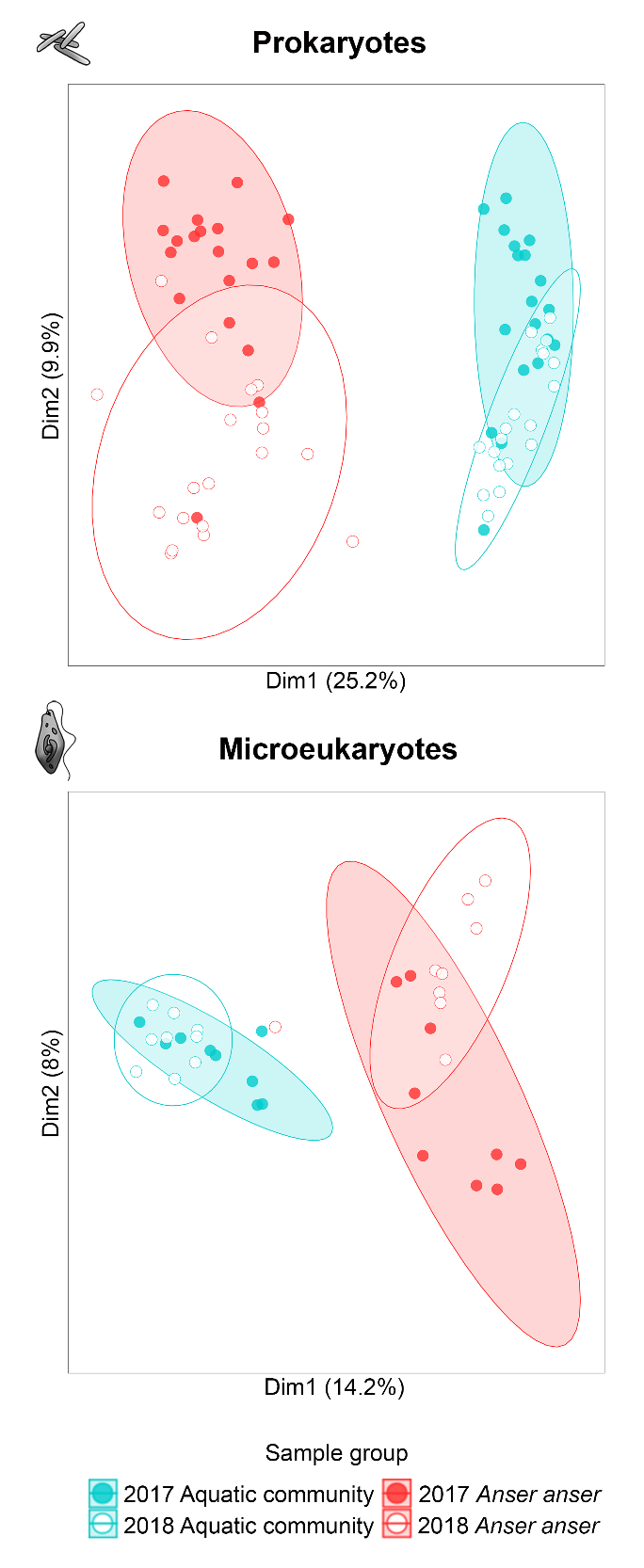

Figure S3. PCoA biplot of aquatic community and *Anser anser* dropping samples collected in 2017 and 2018. The analysis is based on the aquatic subset (incidence data, Sørensen dissimilarity) of prokaryotic and microeukaryotic communities.

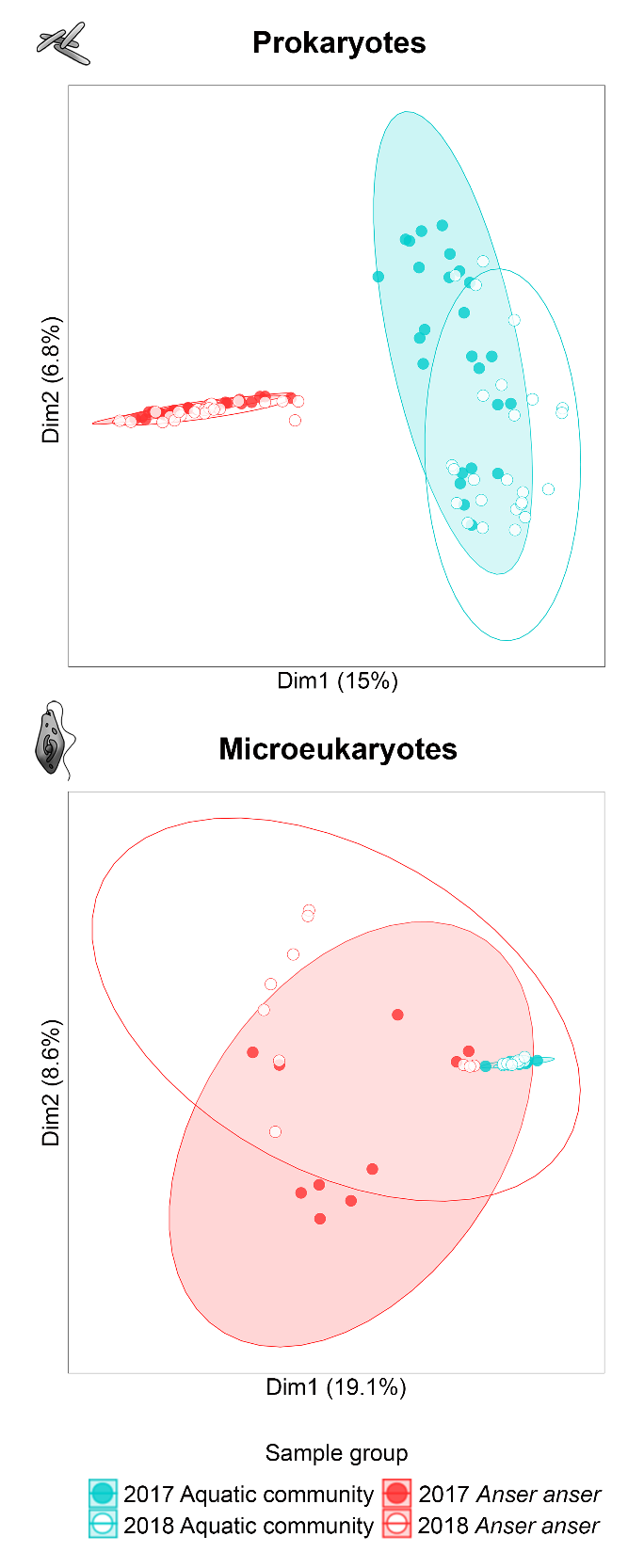

Figure S4. PCoA biplot of aquatic community and *Anser anser* dropping samples collected in 2017 and 2018. The analysis is based on the unselected dataset (abundance data, Bray-Curtis dissimilarity) of prokaryotic and microeukaryotic communities.

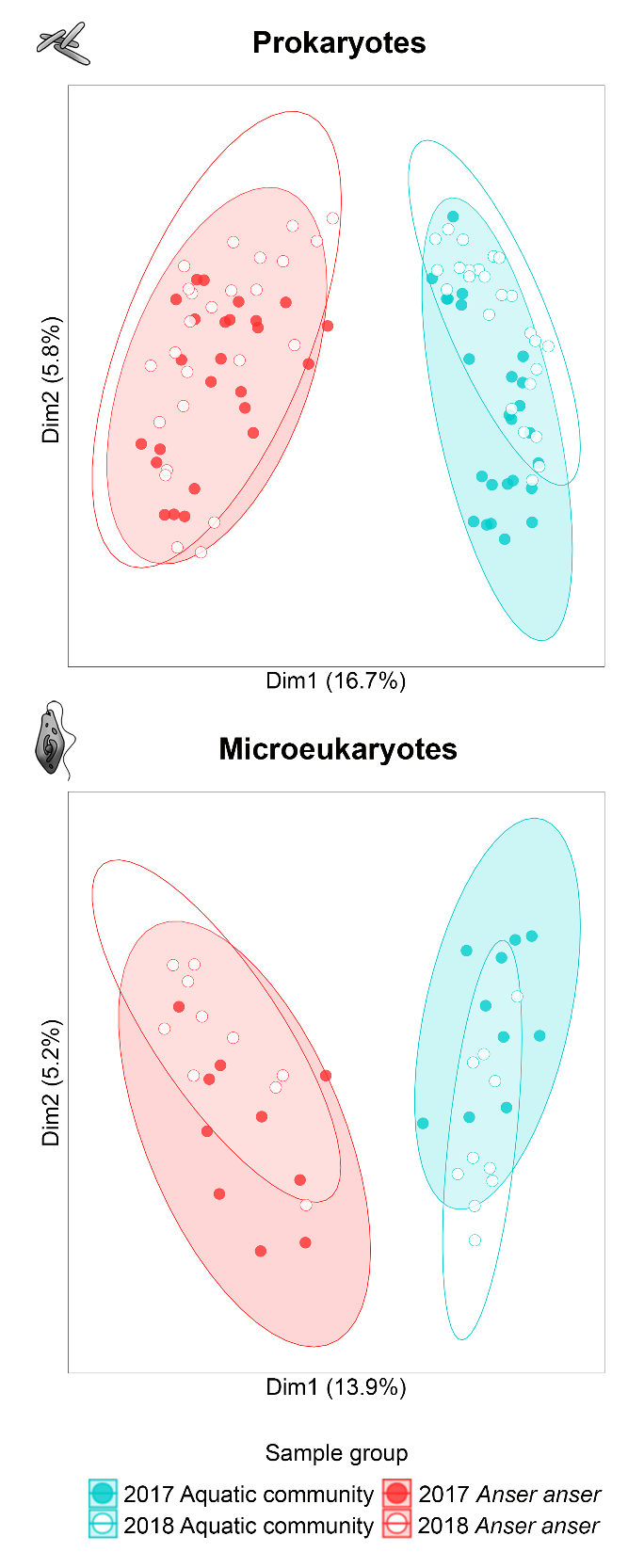

Figure S5. PCoA biplot of aquatic community and *Anser anser* dropping samples collected in 2017 and 2018. The analysis is based on the unselected dataset (incidence data, Sørensen dissimilarity) of prokaryotic and microeukaryotic communities.

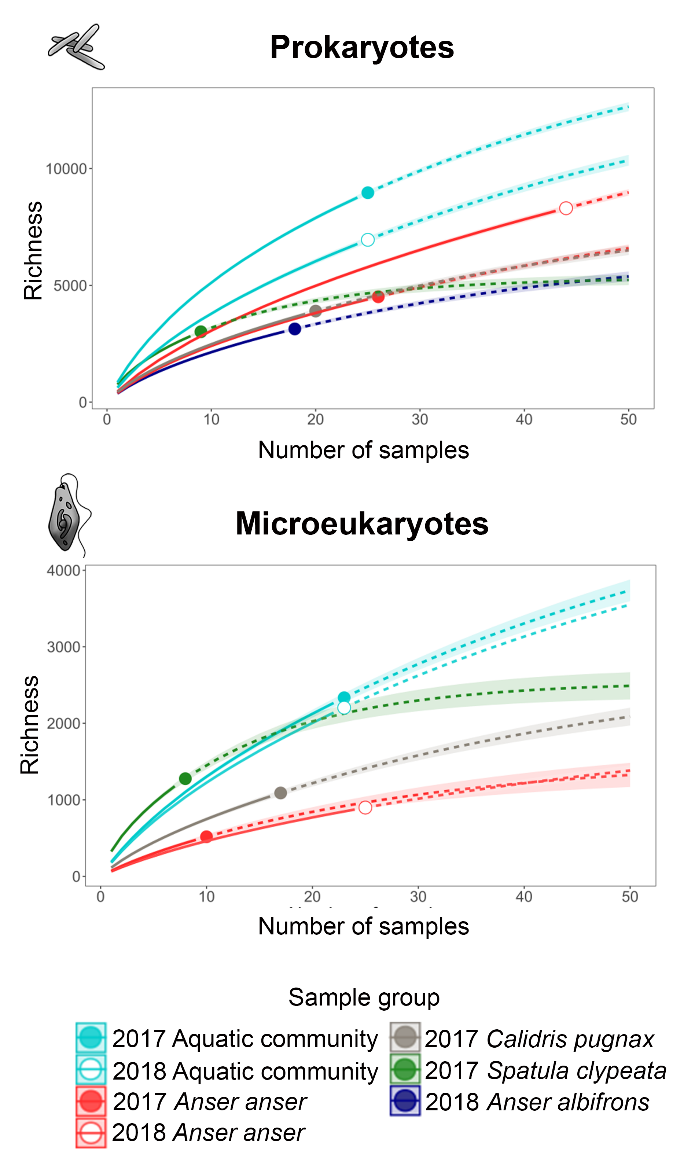

Figure S6. ASV accumulation curves with extrapolated richness estimates and confidence intervals for the unselected datasets of prokaryotes and microeukaryotes dispersed by the five waterbird species compared to the aquatic communities.

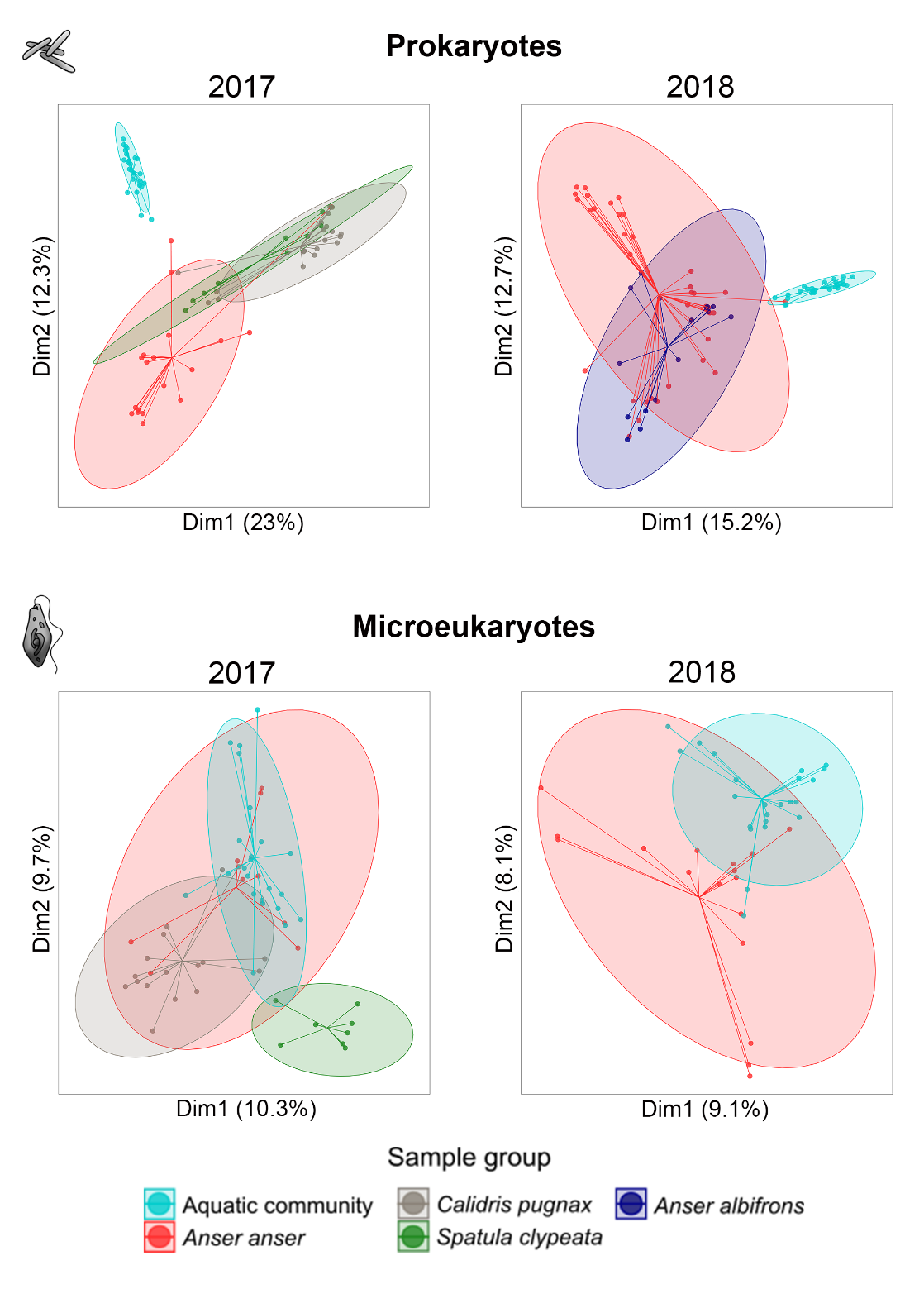

Figure S7. PCoA biplot based on the aquatic subset (abundance data, Bray-Curtis dissimilarity) of prokaryotic and microeukaryotic communities demonstrating the distribution of aquatic community and waterbird dropping samples collected in 2017 and 2018.

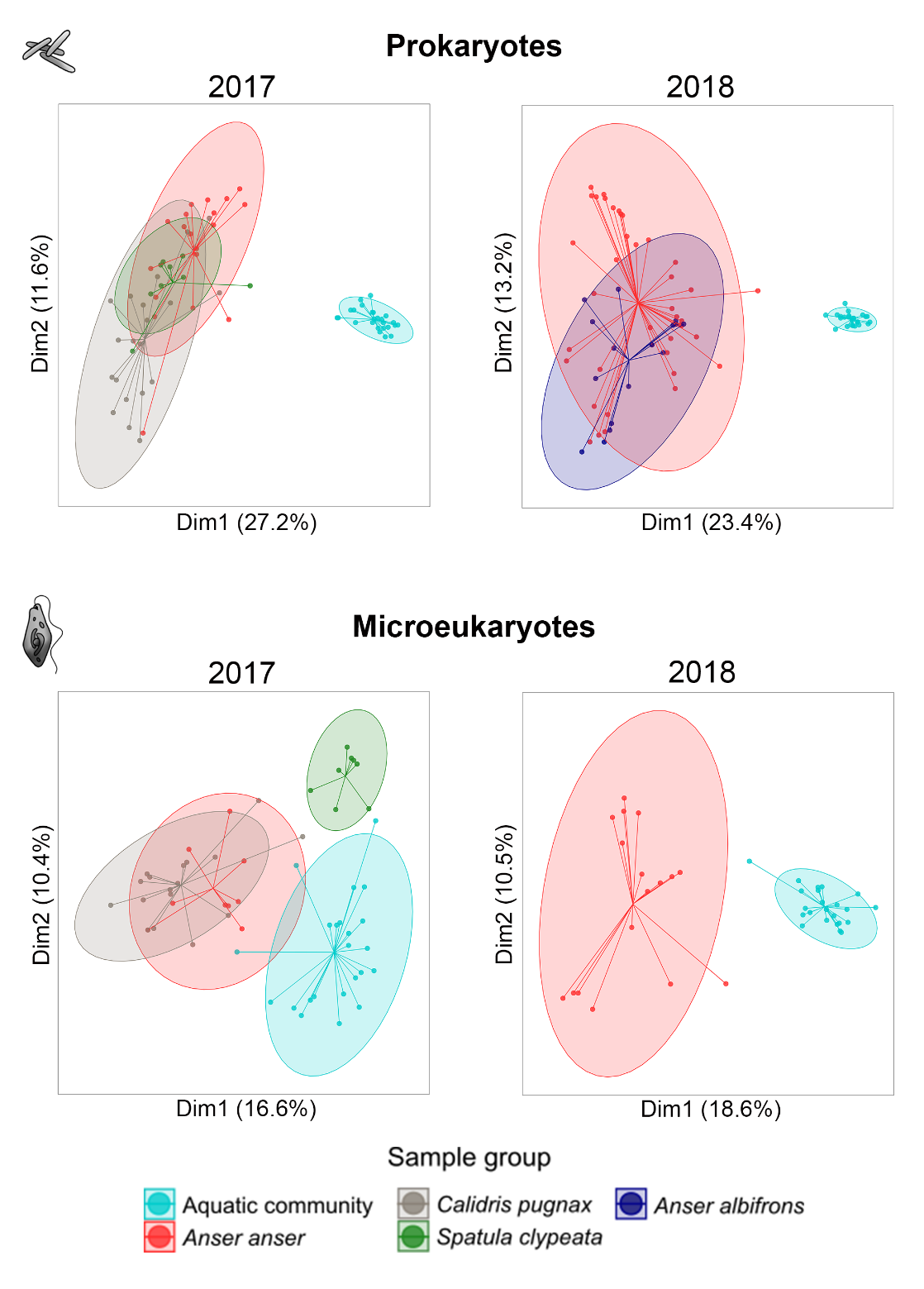

Figure S8. PCoA biplot based on the aquatic subset (incidence data, Sørensen dissimilarity) of prokaryotic and microeukaryotic communities demonstrating the distribution of aquatic community and waterbird dropping samples collected in 2017 and 2018.

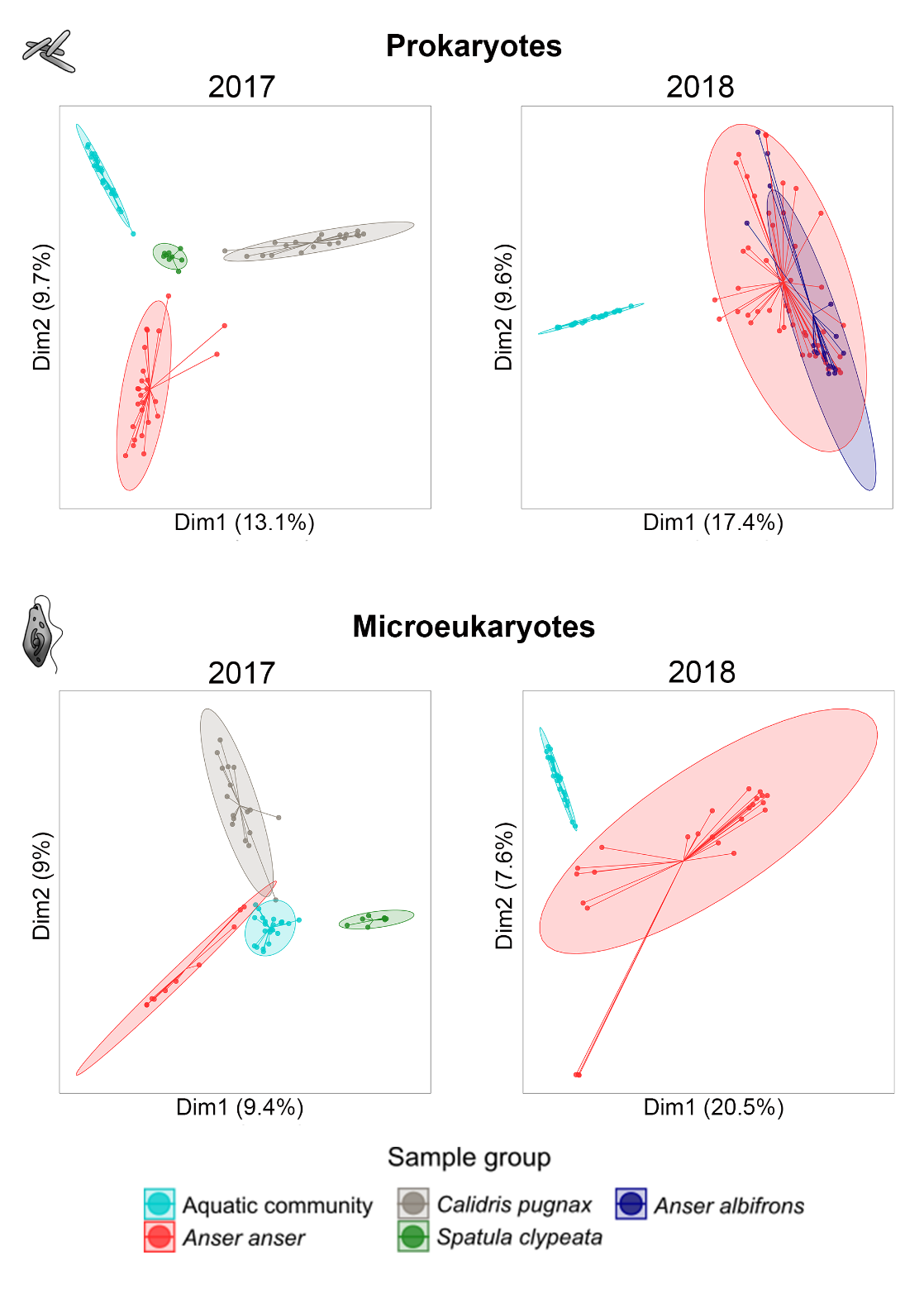

Figure S9. PCoA biplot based on the unselected dataset (abundance data, Bray-Curtis dissimilarity) of prokaryotic and microeukaryotic communities demonstrating the distribution of aquatic community and waterbird dropping samples collected in 2017 and 2018.

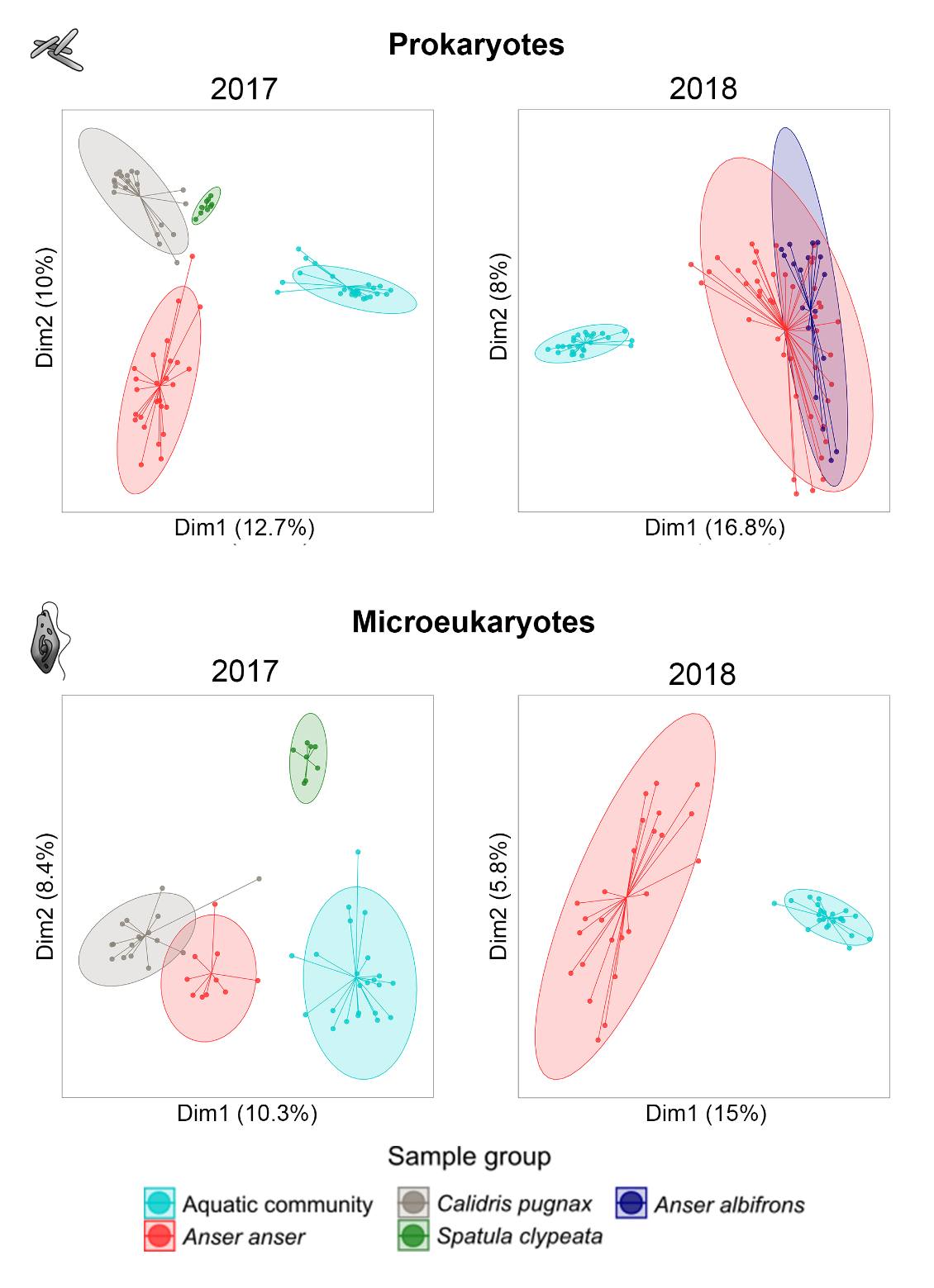

Figure S10. PCoA biplot based on the unselected dataset (incidence data, Sørensen dissimilarity) of prokaryotic and microeukaryotic communities demonstrating the distribution of aquatic community and waterbird dropping samples collected in 2017 and 2018.
